# Resurgent Na^+^ Current Offers Noise Modulation in Bursting Neurons

**DOI:** 10.1101/491324

**Authors:** Sharmila Venugopal, Soju Seki, David H Terman, Antonios Pantazis, Riccardo Olcese, Martina Wiedau-Pazos, Scott H Chandler

## Abstract

Neurons utilize bursts of action potentials as an efficient and reliable way to encode information. It is likely that the intrinsic membrane properties of neurons involved in burst generation may also participate in preserving its temporal features. Here we examined the contribution of the persistent and resurgent components of voltage-gated Na^+^ currents in modulating the burst discharge in sensory neurons. Using mathematical modeling, theory and dynamic-clamp electrophysiology, we show that, distinct from the persistent Na^+^ component which is important for membrane resonance and burst generation, the resurgent Na^+^ can help stabilize burst timing features including the duration and intervals. Moreover, such a physiological role for the resurgent Na^+^ offered noise tolerance and preserved the regularity of burst patterns. Model analysis further predicted a negative feedback loop between the persistent and resurgent gating variables which mediate such gain in burst stability. These results highlight a novel role for the voltage-gated resurgent Na^+^ component in moderating the entropy of burst-encoded neural information.

**Author Summary:** The nervous system extracts meaningful information from natural environments to guide precise behaviors. Sensory neurons encode and relay such complex peripheral information as electrical events, known as action potentials or spikes. The timing intervals between the spikes carry stimulus-relevant information. Therefore, disruption of spike timing by random perturbations can compromise the nervous system function. In this study we investigated whether the widely-distributed voltage-gated sodium (Na^+^) ion channels important for spike generation can also serve as noise modulators in sensory neurons. We developed and utilized mathematical models for the different experimentally inseparable components of a complex Na^+^ channel current. This enabled phenomenological simplification and examination of the individual roles of Na^+^ components in spike timing control. We further utilized real-time closed-loop experiments to validate model predictions, and theoretical analysis to explain experimental outcomes. Using such multifaceted approach, we uncovered a novel role for a resurgent Na^+^ component in enhancing the reliability of spike timing and in noise modulation. Furthermore, our simplified model can be utilized in future computational and experimental studies to better understand the pathological consequences of Na^+^ channelopathies.

## Introduction

Real-time signal detection in uncertain settings is a fundamental problem for information and communication systems. Our nervous system performs the daunting task of extracting meaningful information from natural environments and guides precise behaviors. Sensory neurons for instance use efficient coding schemes such as bursting that aid in information processing^1^. Mathematical models of bursting have helped explain the basic structure of an underlying dynamical system as one in which a slow process dynamically modulates a faster action potential/spike-generating process, leading to stereotypical alternating phases of spiking and quiescence^2, 3^. The so-called *recovery period* of the slow process governs the intervals between bursts which is often susceptible to random perturbations. Uncertainty in spike/burst intervals can alter the timing precision and information in a neural code^4^. Consequently, ionic mechanisms that modulate the recovery of membrane potential during spike/burst intervals, can play a role in maintaining the stability of these timing events and aid neural information processing. Here we examined a candidate mechanism involving neuronal voltage-gated Na^+^ currents for a role in the stabilization of burst discharge (durations and intervals) and noise modulation.

Voltage-gated Na^+^ currents are essential for spike generation in neurons^5^. The molecular and structural diversity of Na^+^ channels and the resultant functional heterogeneity and complexity, suggest their role beyond mere spike generation^6^. For instance, in addition to the fast/transient Na^+^ current (*I*_*NaT*_) mediating action potentials, a subthreshold activated persistent Na^+^ current (*I*_*Nap*_) participates in the generation of subthreshold membrane oscillations (STO) (e.g., see^7^). These oscillations can lead to membrane resonance by which a neuron produces the largest response to oscillatory inputs of some preferred frequency^8, 9^. Neurons utilize this mechanism to amplify *weak* synaptic inputs at resonant frequencies^10^. The slow inactivation and recovery of *I*_*NaP*_ further provides for the slow process required for rhythmic burst generation^11–13^ and therefore can contribute to efficient information processing in multiple ways. However, during ongoing activity, random membrane fluctuations can alter the precision and order of burst timings which can distort/diminish the information in neural code. Here, we provide evidence that a frequently observed resurgent Na^+^ current (*I*_*NaR*_), often coexistent with *I*_*NaP*_ might be a mechanism by which neurons stabilize burst discharge while maintaining its order and entropy. The *I*_*NaR*_, in neurons and other excitable cells is an unconventional Na^+^ current which physiologically activates from a *brief* membrane depolarization followed by repolarization, such as during an action potential ^11, 14–17^ In the well-studied neuronal Nav1.6-type Na^+^ channels, such a macroscopic *I*_*NaR*_ is biophysically suggested to occur from an open-channel block/unblock mechanism^18, 19^. Consequently, *I*_*NaR*_ is known to mediate depolarizing after-potentials and promote high-frequency spike discharge in neurons ^14, 20–24^ Sodium channels containing the Nav1.6 subunits carry all three types of sodium currents and are widely distributed in the central and peripheral neurons and participate in burst generation ^14, 25^. Sodium channelopathy involving alteration in *I*_*NaR*_ and *I*_*NaP*_, and its association with irregular firing patterns and ectopic bursting in disease (e.g., ^26–29^), prompted us to investigate distinct roles for these Na^+^ currents in regulating bursting in sensory neurons.

Lack of suitable functional markers and experimental tools to dissociate the molecular mechanism of *I*_*NaR*_ from *I*_*NaP*_, led us to use computational modeling and dynamic-clamp electrophysiology to examine a role for *I*_*NaR*_ in burst control; however see ^19, 23, 24^. Although existing Markovian models model a single channel Nav1.6 type *I*_*Na*_ using a kinetic scheme (e.g., ^30, 31^), they have limited application for studying exclusive roles of *I*_*NaR*_ and *I*_*NaP*_ in the control of neural bursting; however see^13, 23, 32, 33^. Here, we developed a novel mathematical model for *I*_*NaR*_ using the well-known Hodgkin-Huxley (HH) formalism which closely mimics the unusual voltage-dependent open-channel unblocking mechanism. We integrated the model *I*_*NaR*_ into a bursting neuron model with *I*_*NaP*_ that we previously reported to study their exclusive roles in burst control in the jaw proprioceptive sensory neurons in the brainstem Mesencephalic V (Mes V) nucleus^8^. To validate model predictions, we used in *vitro* dynamic-clamp electrophysiology, and theoretical stability analysis. Using these approaches, we identify a novel role for *I*_*NaR*_ in stabilizing burst discharge and noise modulation in these sensory neurons (see Fig. 1 for a workflow and approaches used).

**Figure 1.**
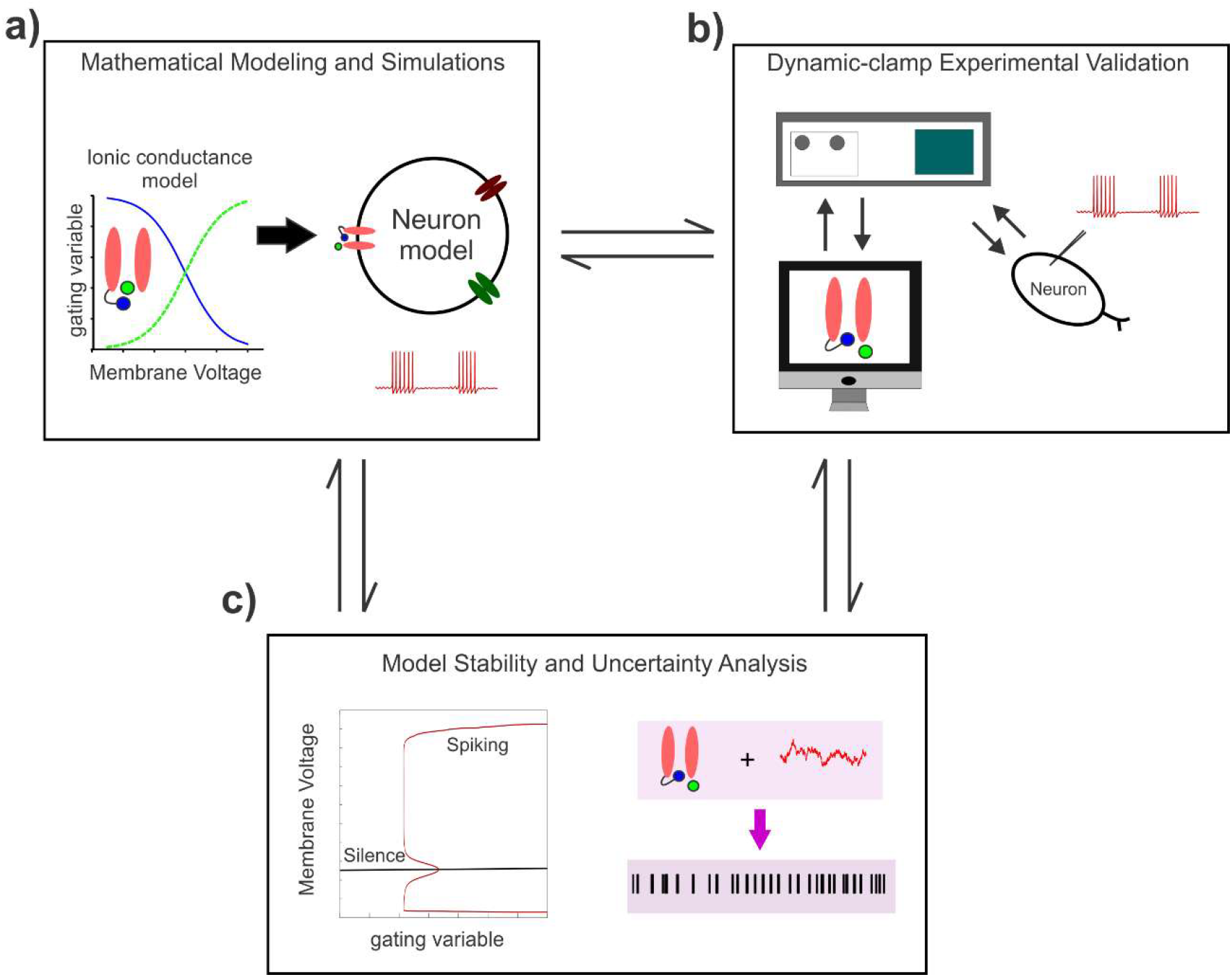
A workflow showing the study components and approaches. **a)** Development of a mathematical model for the Nav1.6-type Na^+^ current components and its incorporation into a minimal conductance-based model for a bursting neuron. **b)** Dynamic-clamp validation of model predictions in the brainstem proprioceptive sensory Mes V neurons. **c)** The observed effects on neural discharge are explained using theoretical stability and uncertainty analyses.

## Results

We first developed a novel HH-based model for the resurgent component of a Nav1.6-type Na^+^ current and combined it with our previous HH-type models for the transient and persistent Na^+^ currents^8^. Although the total Na^+^ current arises from a single channel^11, 30^, we formulated the model *I*_*Na*_ as a sum of the three components as shown in Fig. 2a (also see **Methods**): the transient, *I*_*NaT*_, the resurgent, *I*_*NaR*_, and, the persistent, *I*_*NaP*_ current. This enabled easy manipulation of individual components to study their exclusive role in burst control. We ensured that each of these components mimic the distinct voltage-dependencies and kinetics observed during voltage-clamp recording in Mes V neurons^11, 34^ The classic *I*_*NaT*_ with fast inactivation kinetics (order of 1 ms) mediates spike generation, the persistent *I*_*NaP*_ with slow inactivation/recovery (order of 1000 ms) mediates STO and provides for the slow process responsible for rhythmic bursting in Mes V neurons^8, 35, 36^, and, the resurgent *I*_*NaP*_ which mimics the phenomenology of an open channel unblock mechanism with a peak time of ~6 ms and decay kinetics on the order of 10 ms^11, 19, 30^ (also see Fig. 3).

**Figure 2.**
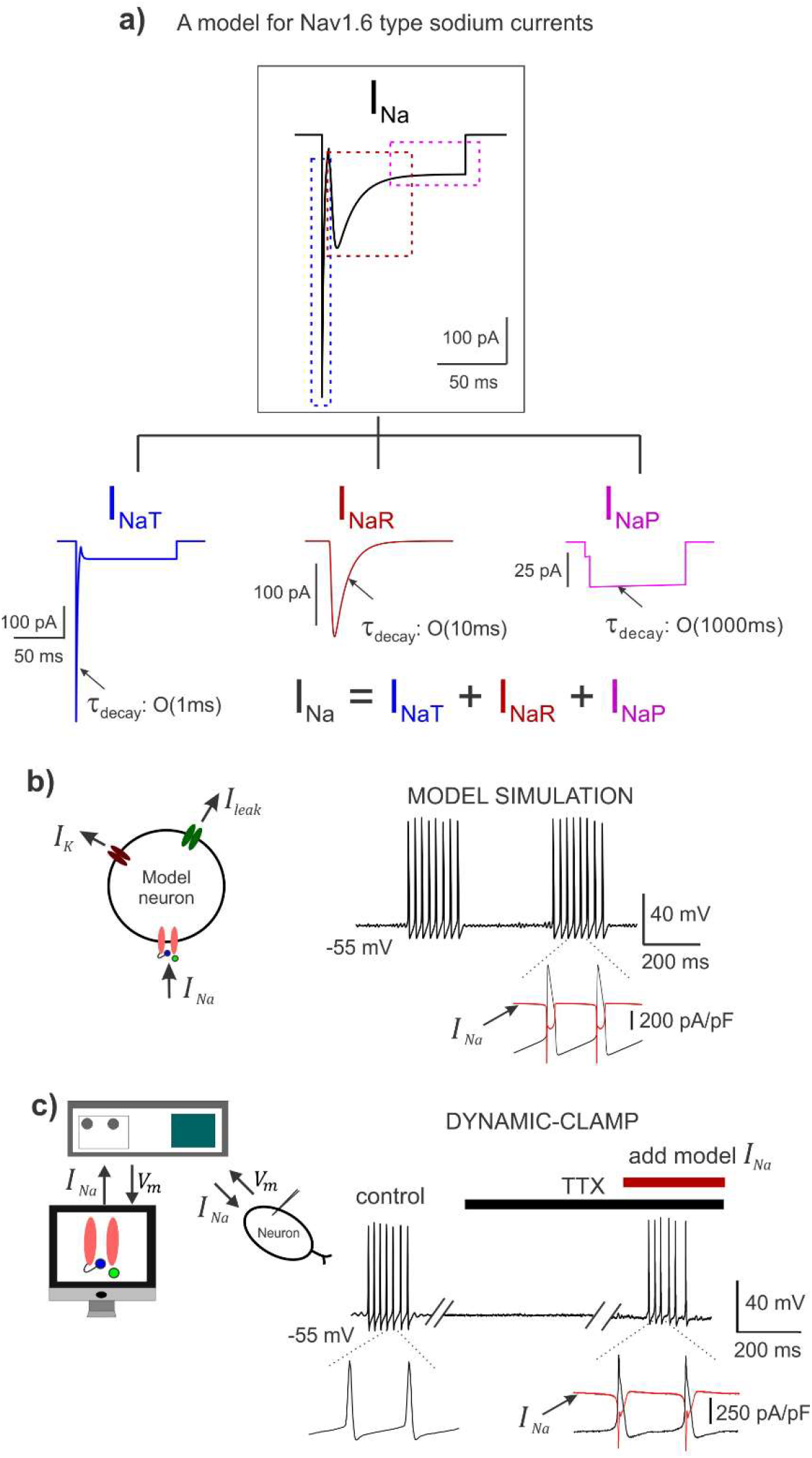
A Hodgkin-Huxley model for the Nav1.6 Na^+^ current with the three components. **a)** A simulated trace showing total *I*_*Na*_; the tree map shows each of the three components: transient *I*_*NaT*_, resurgent *I*_*NaR*_, and, persistent *I*_*NaP*_; These are also highlighted by the color-matched dashed boxes in the top panel. The *τ*_*decay*_ shows the order of magnitude of the decay kinetics of the three components. **b)** *Left:* A schematic showing a conductance-based minimal model for a bursting neuron with the *I*_*Na*_ incorporated; other ionic currents in the model include a delayed-rectifier potassium (*I*_*K*_) and a leak current (*I*_*leak*_). *Right:* Model simulation demonstrates rhythmic burst discharge and inset highlights the *I*_*Na*_ current in the model during action potentials, in red. **c)** *Left:* Schematic shows the dynamic-clamp experimental approach; *V*_*m*_ is the membrane potential. *Right*: Membrane potential recorded from a rhythmically bursting sensory neuron; action potentials were blocked using 1*μ*M TTX, and dynamic-clamp model *I*_*Na*_ was applied to regenerate spikes; double slanted lines indicate break in time; inset highlights the dynamic-clamp *I*_*Na*_ in red, during action potentials.

**Figure 3.**
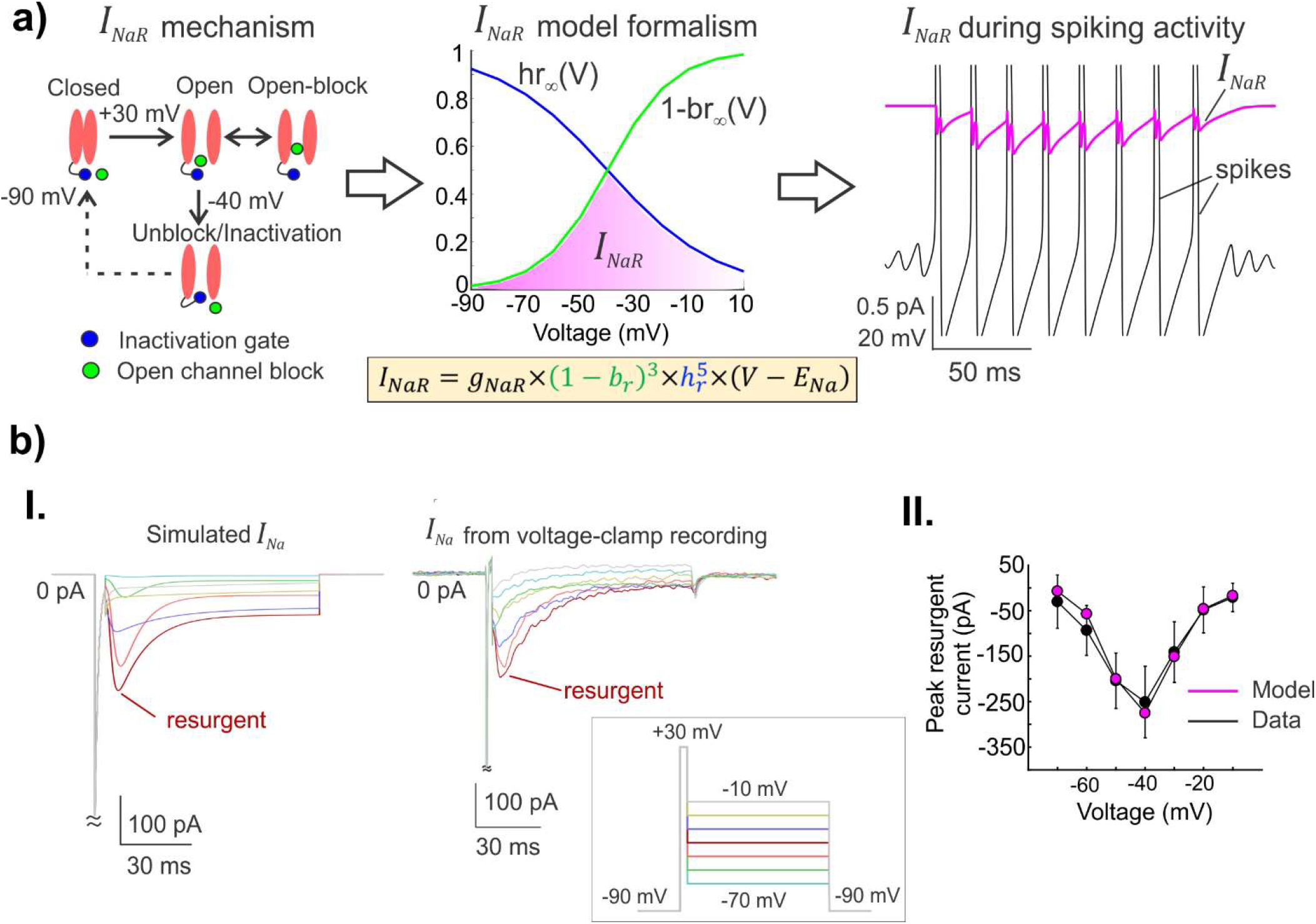
A novel mathematical model for the unusual resurgent component of the Nav1.6-type Na^+^ current. **a)** *Left:* A schematic showing the voltage-dependent operation of a Na+ channel mediating resurgent current; open-channel block/unblock (green circle); classic inactivation gate (blue ball and chain). *Middle:* The steady-state voltage-dependencies of open-channel unblocking (1 – *br*_∞_(*V*)) and a competing inactivation process *hr*_∞_(*V*) for the novel resurgent component are shown (also see Methods and Results); the magenta shaded region highlights the voltage range over which *I*_*NaR*_ can be observed during open-channel unblocking. The equation for *I*_*NaR*_ is shown with novel blocking (*b*_*r*_), and inactivation (*h*_*r*_) gating variables. *Right:* The simulated *I*_*NaR*_ (in magenta) with peaks occurring during the repolarization phase of action potentials (in black) is shown. **b) I.** A comparison between simulated *I*_*Na*_ and experimentally generated *I*_*Na*_ from voltage-clamp recording is shown; boxed inset shows the experimental protocol typically used to test for voltage-dependent activation of *I*_*NaR*_. **II.** Graphs show the nonlinear current-voltage relationship of peak resurgent current in the model (magenta) and average peak resurgent currents measured from voltage-clamp experiments (black); error bars show standard deviation (*n* = 5 neurons from 5 animals).

We compared our model *I*_*Na*_ and its components to the macroscopic *I*_*Na*_ observed in voltage-clamp recordings in Mes V neurons^11, 14^ to ensure qualitative and quantitative similarities (see Fig. 3b). We further ensured that our *I*_*Na*_ model closely resembled the *I*_*Na*_ generated in Markovian models^30^, which follow a kinetic scheme and do not formulate the three components separately (see **Supplementary Fig. 1**). Our model *I*_*Na*_ also satisfied previously described contingency of Nav1.6-type Na^+^ channels carrying resurgent current: activation of *I*_*NaP*_ only up on brief depolarization followed by hyperpolarization to ~ - 40 mV, and the correspondence between maximum channel availability and peak time of resurgent current^30^ (see **Supplementary Fig. 2**).

We incorporated the model *I*_*Na*_ into a conductance-based single-compartment neuron model (see schematic and membrane voltage trace in black in Fig. 2b). Together with a minimal set of Na^+^, K^+^ and leak conductances, the model neuron faithfully reproduced an expected rhythmic burst discharge observed in Mes V sensory neurons; the total *I*_*Na*_ generated during action potentials is shown in expanded time in the figure (red trace). Figure. 2c shows the dynamic-clamp experiment in an intrinsically bursting Mes V neuron. The control burst was generated by simply driving the neuron with a step depolarization, following which we blocked action potential generation by bath application of tetrodotoxin (1 *μ*M TTX) (see black horizontal bar in Fig. 2c). Subsequently, we introduced the real-time model *I*_*Na*_ using dynamic-clamp during TTX application and by adjusting the conductances of the three *I*_*Na*_ components suitably, we were able to regenerate action potential bursts (see **Methods** on the choice of conductance values). The dynamic-clamp *I*_*Na*_ generated during action potentials is shown in expanded time in the figure for comparison with the model simulation in Fig. 2b (red trace).

### A novel HH-based mathematical model for resurgent Na^+^ current

As noted earlier, the total *I*_*Na*_ in our model has the novel resurgent component, *I*_*NaR*_; the transient and persistent components are similar to our previous report^8^. Figure 3a **(left panel)** illustrates a well-accepted mechanism of Na^+^ resurgence^30^, wherein a putative blocking particle occludes an open channel following brief depolarization such as during an action potential; subsequently upon repolarization, a voltage dependent unblock results in a resurgent Na^+^ current. Our *I*_*NaR*_ formulation recapitulates this unusual behavior of Na^+^ channels using nonlinear ordinary differential equations for a blocking variable (*b*_*r*_) and a competing inactivation (*h*_*r*_) (see **Methods**). Different from a traditional activation variable of an ionic current in HH models, we formulated the *I*_*NaR*_ gating using a term (1 − *b*_*r*_) to enable an unblocking process (see *I*_*NaR*_ equation in **Methods** and in Fig. 3a). Here, the model variable *b*_*r*_ reflects the fraction of channels in the blocked state at any instant, and ‘1’ denotes the maximum proportion of open channels, such that (1 – *b*_*r*_) represented the fraction of unblocking channels. Such a formulation enabled mimicking the open-channel unblocking process as follows: Normally the *b*_*r*_ is maintained ‘high’ such that (*1 – b_r_*) term is small and therefore no *I*_*NaR*_ flows, except when a spike occurs, which causes *b*_*r*_ to decay in a voltage-dependent manner. This turns on the *I*_*NaR*_ which peaks as the membrane voltage repolarizes to ~ −40 mV, following which the increasing inactivation variable *h*_*r*_ gradually turns off the *I*_*NaR*_. The steady-state voltage dependency of unblock, (l – *br*_*∞*_(*V*)), and the competing inactivation (*hr*_*∞*_ (*V*)) in the model are shown in Fig. 3a **(middle panel)**, along with the equation for *I*_*NaR*_; the magenta shaded region highlights the voltage-dependency of *I*_*NaR*_ activation as posited to occur during open-channel unblocking. In Fig. 3a **(right panel)**, we show simulated *I*_*NaR*_ (in magenta), peaking during the recovery phase of spikes (in black). In Fig. 3b **(I.)**, we reproduced experimentally observed *I*_*Na*_ during voltage-clamp recording and highlight the resurgent component in both model *(left)* and experiment *(right*), (inset shows experimental protocol; also see legend and **Methods**). A comparative current-voltage relationship for the model and experiments is shown in Fig. 3b **(II.)**; also see **Supplementary Fig. 3** for detailed kinetics of model *I*_*NaR*_. Taken together, the above tests and comparisons ensured the suitability of our model for further investigation of *I*_*NaR*_ mediated burst control.

### Resurgent and persistent Na^+^ currents offer *push-pull* modulation of spike/burst intervals

Given that *I*_*NaR*_ is activated during the recovery phase of an action potential, physiologically, any resulting rebound depolarization may control the spike refractory period, and increase spike frequency and burst duration^22, 37^. We tested this by selectively increasing the maximal resurgent conductance *g*_*NaR*_ in our model neuron simulation and validated the predictions using dynamic-clamp experiments as shown in Figs. 4a, b. In parallel, we also exclusively modified the maximal persistent conductance *g*_*Nap*_ using model simulations and verified the effects using dynamic-clamp experiments as shown in Figs. 4c, d. We only focused on rhythmically bursting Mes V neurons and quantified the burst timing features including the inter-burst intervals (IBIs), burst duration (BD) and inter-spike intervals (ISIs) as illustrated in the boxed inset in Fig. 4. **Figure panels 4e - j** show a comparison of the exclusive effects of *g*_*NaR*_ versus *g*_*NaP*_ on each of these burst features. These experimental manipulations using model currents and quantification of resulting burst features revealed significant differences and some similarity between the action of *g*_*NaR*_ and *g*_*NaP*_ in burst control. First, increases in *g*_*NaR*_ resulted in *longer* IBIs, which was in contrast with the effects of the persistent Na^+^ conductance, *g*_*Nap*_, which *decreased* IBIs (effect highlighted with red double arrows in Figs. 4a - d and quantified using box plots in Figs. 4e and h). Alternatively, increasing *g*_*NaR*_ *reduced* ISIs, while *g*_*NaP*_ had the opposite effect on these events, resulting in an overall *increase* in ISIs with *g*_*NaP*_ increases (see Figs. 4f and i); whereas *g*_*NaR*_ and *g*_*NaP*_ had similar effect in increasing BDs (see Figs. 4g and j). Box plots in Figs. 4e - j show 1^st^, 2^nd^ (median) and 3^rd^ quartiles; error bars show 1.5x deviations from the inter-quartile intervals. For each case (or cell), *n* values represented events (ISI, IBI and BD) and reported here are within-cell effects for different *g*_*NaR*_ and *g*_*NaP*_ applications during 20 sec step-current stimulation. For the *g*_*NaR*_ applications (or series), IBI mean ± std were 210.49 ± 69.33 (control, *n* = 7), 450.29 ± 116.56 (1x, *n* = 26) and 1074.06 ± 199.17 (2x, *n* = 11), and for *g*_*NaP*_ applications, these values were 1448.71 ± 450.92 (control, *n* = 4), 910.10 ± 527.27 (1x, *n* = 8) and 644.35 ± 234.19 (1.5x, *n* = 6). For *g*_*NaR*_ series, ISI mean ± std were 20.26 ± 1.65 (control, *n* = 59), 12.56 ± 2.25 (1x, *n* = 646) and 10.56 ± 1.81 (2x, *n* = 783), and for *g*_*NaP*_ series these values were 12.76 ± 1.13 (control, *n* = 159), 12.57 ± 1.05 (1x, *n* = 611) and 13.56 ± 1.42 (1.5x, *n* = 503). For g_*NaR*_ series, BD mean ± std were 176.36 ± 69.57 (control, *n* = 8), 300.56 ± 77.90 (1x, *n* = 27) and 671.16 ± 285.96 (2x, *n* = 12), and for *g*_*NaP*_ series these values were 346.59 ± 88.39 (control, *n* = 4), 571.30 ± 197.56 (1x, *n* = 8) and 1334.20 ± 592.59 (1.5x, *n* = 7). Treatment effects and group statistics for all the replicates showing the above effects across six Mes V bursting neurons, each from a different animal are summarized in **Supplementary Table 2**. A one-way ANOVA was used to test the treatment effects of *g*_*NaR*_ and *g*_*NaP*_ applications, and when significant, a *post hoc* two-sample Student t-test was used for group comparisons between control and 1x, 1x and 2x, and control and 2x for *g*_*NaR*_ cases, and between control and 1x, 1x and 1.5x, and control and 1.5x for *g*_*NaP*_ cases. Asterisks above box plots between groups in Figs. 4e - j indicate *p* < 0.05 using two-sample Student t-tests for *post-hoc* comparisons. Furthermore, as shown in Fig. 4, the model simulations predicted consistent effects (note white circles denote predicted values in panels **e - j**) with dynamic-clamp experiments, making the model suitable for further analysis of *I*_*NaR*_/*I*_*NaP*_ mediated mechanism of burst control (see *white circles* in panels **4e - j**). In two additional bursting neurons, we also conducted *g*_*NaR*_ and *g*_*Nap*_ subtraction experiments which showed consistent reverse effects of additions (see **Supplementary Fig. 4 and legend**). In the *g*_*Nap*_ subtraction experiment, note that a −2x *g*_*NaP*_ resulted in the abolition of bursting and sub-threshold oscillations as shown in the figure. Such an effect was reproduced in the neuron model by setting *g*_*NaP*_ = 0, as shown in **Suppl. Fig. 4c**.

**Figure 4.**
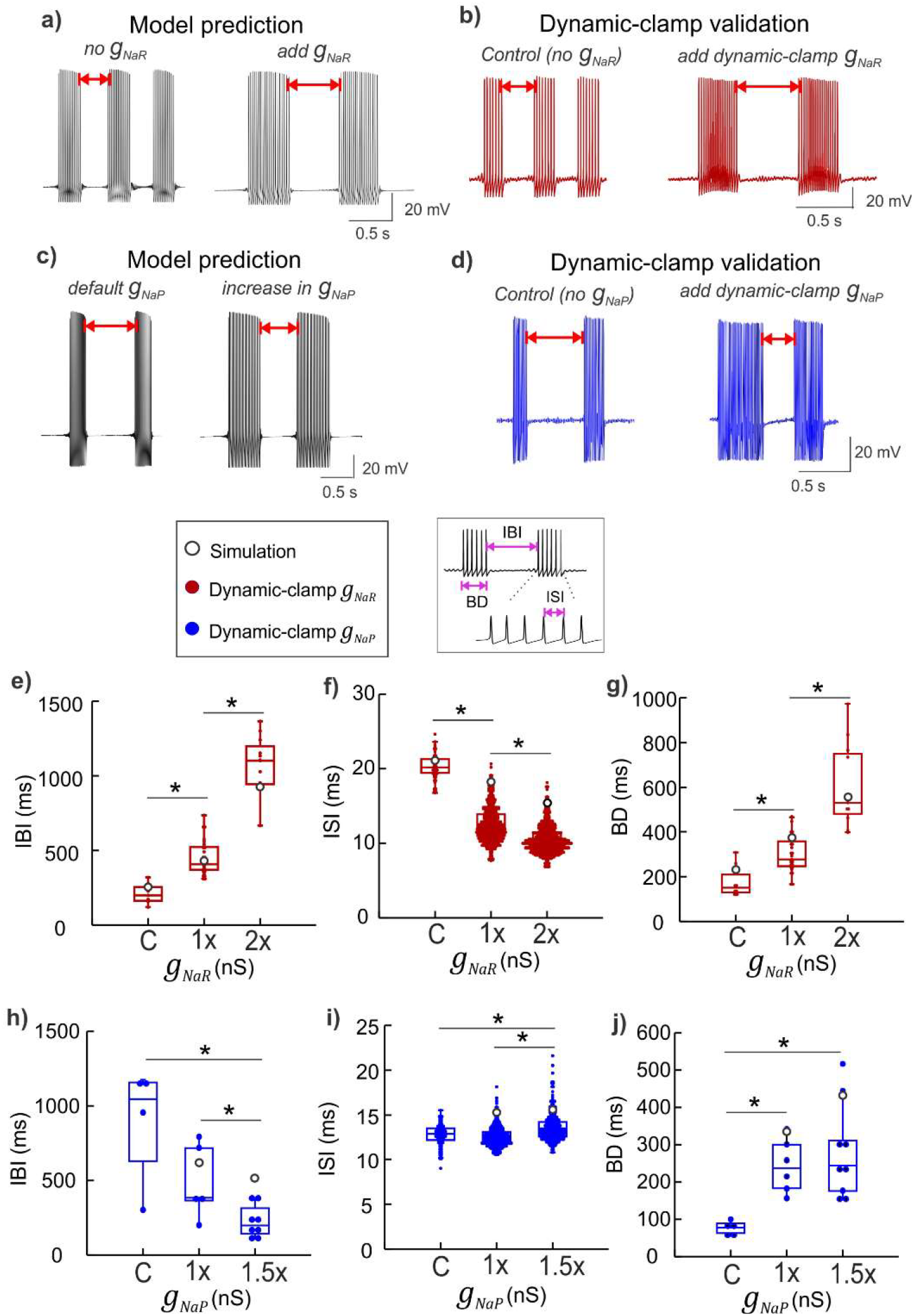
Physiological consequences of *I*_*NaR*_ and *I*_*NaP*_ on burst discharge. **a, c)** Simulated membrane voltage demonstrating the effect of *g*_*NaR*_ addition **(a)**, and *g*_*NaP*_ increase **(c)** on burst discharge. **b, d)** Dynamic-clamp experimental traces showing effects of real-time addition of *g*_*NaR*_ **(b)** and *g*_*NaP*_ **(c)** in intrinsically bursting sensory neurons; the red double arrows highlight the opposite effects of increases in *g*_*NaR*_ and *g*_*NaP*_ on inter-burst intervals in panels **(a - d)**. The boxed inset *(left)* indicates color coding of traces for panels **e - j.** The boxed inset *(right)* highlights burst features quantified in panels **(e - j)**: inter-burst intervals (IBI), burst duration (BD), and inter-spike intervals (ISI. **e - j)** Box plots showing IBIs **(e, h)**, BDs **(f, i)**, and ISIs **(g, j)** for application of a series of *g*_*NaR*_ or *g*_*NaP*_ conductance values (see Results for statistics). Model values of 1x *g*_*NaR*_ and *g*_*NaP*_ are adjusted to match experimental data for 1x application of the corresponding conductance tested; C is control, values of *g*_*NaR*_ used for dynamic-clamp are 2 and 4 nS/pF for 1x and 2x respectively, and, values of *g*_*NaP*_ used are 0.25 and 0.375 nS/pF for 1x and 1.5x respectively. Asterisks indicate a *p* < 0.05 using a Student t-Test for group comparisons.

To test whether *I*_*NaR*_ and *I*_*NaP*_ differentially modulate the regularity and precision of spike timing we observed the inter-event intervals (or IEIs) during real-time addition of *g*_*NaR*_ and *g*_*NaP*_. As shown in Fig. 5a, b addition of *g*_*NaR*_ improved the regularity of the two types of events: the longer IBIs and the shorter ISIs, whereas, addition of *g*_*NaP*_ did not show such an effect. Note that application of real-time *g*_*NaR*_ improved both the timing precision and regularity of IBIs (compare *left* and *right* panels of Fig. 5a), while application of dynamic-clamp *g*_*NaP*_ did not seem to either of these features (compare *left* and *right* panels of Fig. 5b). Furthermore, in Fig. 5c we highlight that *g*_*NaR*_ application also improved spike-to-spike regularity of ISIs within bursts (shown are two representative bursts for *g*_*NaR*_ (Fig. 5c, *left* panel) versus *g*_*NaP*_ (Fig. 5c, *right* panel) application from Fig. 5a, b (see figure legend). Taken together, these opposite effects of persistent and resurgent Na^+^ currents arising from a single Na^+^ channel may act in concert to offer a *push-pull* modulation of burst timing regulation.

**Figure 5.**
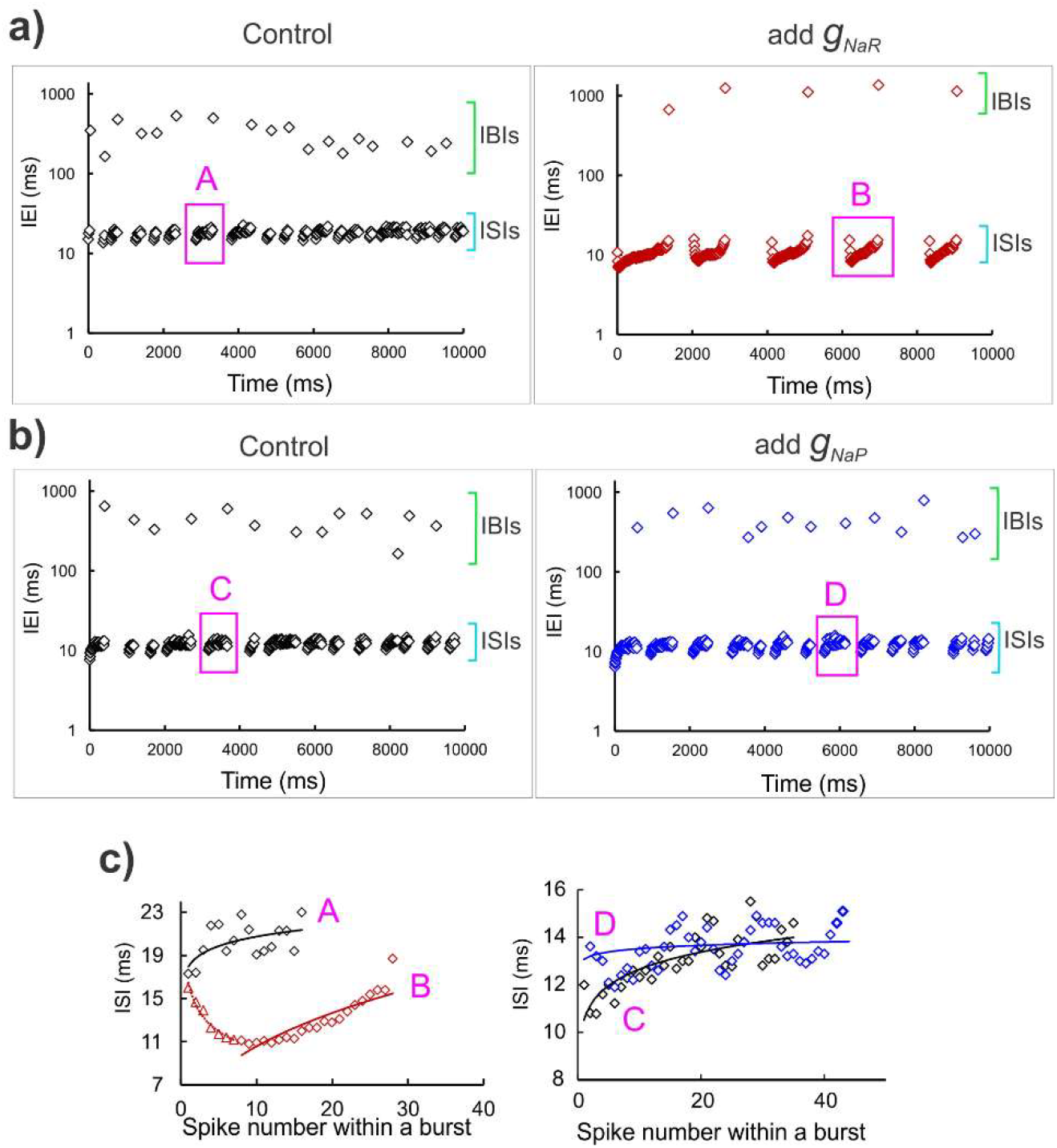
Spike timing regularity monitored by *I*_*NaR*_. **a, b)** Time series graphs showing inter-event intervals (IEIs) under control condition *(left panels* in both **a, b**), and dynamic-clamp addition of *g*_*NaR*_ (*right panel* in **a**), and *g*_*NaP*_ (*right panel* in **b**). The green and cyan square brackets in each panel highlight the IBI and ISI range respectively. The y-axis shows IEIs on a log scale. Magenta boxes demarcate a representative burst in each case, which is expanded in the graphs in **c**), corresponding with the bold letters, A, B, C and D. The solid lines in **(c)** are power regression fit in each case; in all the panels, black open diamonds are control, maroon open diamonds correspond with *g*_*NaR*_ application and blue open diamonds correspond with the *g*_*NaP*_ application.

### Resurgent Na^+^ aids stabilization of burst discharge

The effect of persistent *g*_*NaP*_ in reducing IBIs can be explained by its sub-threshold activation^11^, wherein increases in *g*_*NaP*_ *promotes* STO and burst initiation which in turn increases burst frequency and reduces IBIs. Additionally, a high *g*_*NaP*_ together with its slow inactivation helps maintain depolarization which can increase BDs. During spiking, *I*_*NaP*_ inactivation slowly accumulates and can contribute to any spike frequency adaptation as a burst terminate. Further increases in *g*_*NaP*_ can accentuate such an effect and lead to increases in spike intervals within a burst. In contrast, the rebound depolarization produced due to *I*_*NaR*_ can decrease ISIs and promote spiking which prolongs the BDs. An effect which is not immediately obvious is an increase in IBI due to increases in *g*_*NaR*_ since IBIs are order of magnitude slower events than rise/decay kinetics of *I*_*NaR*_. Moreover, this current is not active during IBIs. It is likely that *I*_*NaR*_ which promotes spike generation produces its effect on lengthening the IBIs by further accumulation of *I*_*NaP*_ inactivation during spiking and in turn reducing channel availability immediately following a burst. We examined this possibility using model analysis of *I*_*NaR*_’s effect on modulating the slow *I*_*NaP*_ inactivation/recovery variable, *h*_*p*_.

The simulated membrane potential (grey traces) and the slow *I*_*NaP*_ inactivation/recovery variable, *h*_*p*_ (overlaid magenta traces) of the model neuron under three conditions shown in Figs. 6a-c: 1) with default values of *g*_*NaR*_ and *g*_*NaP*_ (Fig. 6a), 2) an 1.5x increase in *g*_*NaP*_ (Fig. 6b), and, 3) a 2x increase in *g*_*NaR*_ (Fig. 6c). The peak and trough of the slow inactivation/recovery variable, *h*_*p*_ correspond to burst *onset* and *offset* respectively. Comparing these traces in the three panels, we note that an increase in *g*_*NaR*_ effectively lowers the *h*_*p*_ value at which burst terminates (see curvy arrow in Fig. 6c and legend). This observation was further supported by estimations of theoretical thresholds for *burst onset* and *offset* for increasing values of *g*_*NaR*_ (Fig. 6d) and, similar thresholds for increasing values of *g*_*NaP*_ are provided for comparison in Fig. 6e (see legend and **Supplementary Information** for details). Note that changes in *g*_*NaR*_ did not alter the burst onset thresholds, consistent with a lack of resurgent current before spike onset (see brown arrow indicating burst onset threshold in Fig. 6d). In contrast, increasing *g*_*NaR*_, consistently *lowered* the threshold values of slow inactivation/recovery for burst offset (see highlighted dashed box with arrows pointing to the burst offset thresholds decreasing with increasing *g*_*NaR*_ values in Fig.6d). The net effect is longer recovery time between bursts and therefore prolonged IBIs. Additionally, in Fig. 6d, an increase in *g*_*NaR*_ extended the range of slow inactivation/recovery for which stable spiking regime exists (marked by the green circles). This *gain in stability* is indicative of a *negative feedback loop* in the Na^+^ current gating variables which model the slow inactivation and the open-channel unblock process. As shown in Fig. 6f (see boxed inset), during a burst, presence of a channel unblocking process and the resulting resurgent Na^+^ can lead to further accumulation of slow inactivation (a positive effect) which eventually shuts off the unblocking events with further inactivation (negative feedback), which terminates a burst. The schematic on the left in Fig. 6f summarizes a negative feedback loop between the unblocking and slow inactivation processes of Na^+^ currents which could mediate the stabilization of burst discharge as described above.

**Figure 6.**
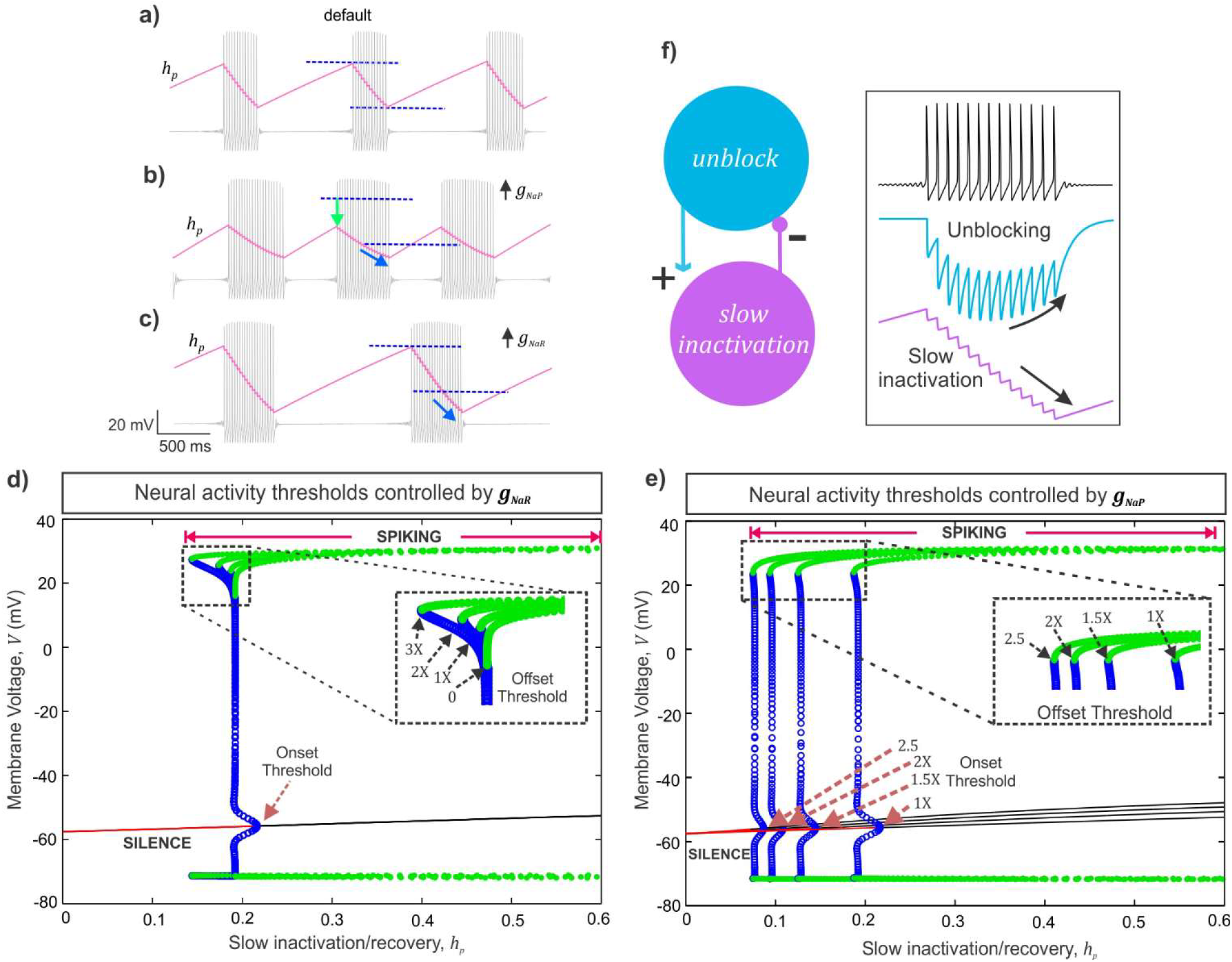
Analysis of a mechanism of action of *I*_*NaR*_ and *I*_*Nap*_. **a-c)** The slow *I*_*NaP*_ inactivation/recovery variable, *h*_*P*_ is overlaid (magenta) on membrane voltage traces (grey) under default **(a)**, 1.5x *g*_*NaP*_ **(b)**, and 2x *g*_*NaR*_ **(c)** conditions; the dark blue dashed lines in **(a-c)** indicate the maximum and minimum values of the persistent inactivation variable under default condition; the light green arrow in **(b)**, highlights a reduced peak recovery required for burst onset; the light blue curvy arrows in **(b, c)** indicate further accumulation of slow inactivation before burst termination. **d, e**) Bifurcation diagrams showing the steady states and spiking regimes of the membrane voltage (*V*) and slow *I*_*NaP*_ inactivation/recovery (*h*_*p*_). The red solid lines represent resting/quiescence states consistent with low values of *h*_*p*_. The meeting points of stable equilibria (red) and unstable equilibria (black solid lines) represents the theoretical threshold for burst onset, the Hopf bifurcation point (see **Supplementary Text**). The blue open circles are the unstable periodics that form region of attraction on either side of the stable equilibria for sub-threshold membrane voltage oscillations; the meeting point of the curve of unstable periodics with the stable periodics (green filled circles) represents the theoretical threshold for burst offset/termination, saddle node of periodics (see Supplementary Text). The dashed boxes in **(d)** and **(e)** magnify the burst offset thresholds due to increases in *g*_*NaR*_ **(d)** and *g*_*Nap*_ **(e)**; brown dashed arrows in **(e)** highlight shifts in burst onset thresholds due to *g*_*NaP*_ increases; 1X *g*_*NaR*_ = 3.3 nS/pF and 1X *g*_*NaP*_ = 0.5 nS/pF.

### Resurgent Na^+^ current offers noise tolerance

Next, we examined whether the stability of burst discharge offered by the presence of *I*_*NaR*_ can also contribute to noise tolerance. During quiescence/recovery periods between bursts, the membrane voltage is vulnerable to perturbations by stochastic influences, which can induce abrupt spikes and therefore disrupt the burst timing precision. We introduced a Gaussian noise to disrupt the rhythmic burst discharge in the model neuron when no *g*_*NaR*_ was present as shown in Figs. 7a, b. Subsequent addition of *g*_*NaR*_ restored burst regularity (Fig. 7c). Model analyses in the (*h*_*p*_, *V*) plane provides insight into the mechanism of *I*_*NaR*_ mediated noise tolerance. Briefly, we project a portion of a (*h*_*p*_,*V*) trajectory corresponding to the termination of one burst until the beginning of the next (see expanded insets in Figs. 7a, b, c) to the (*h*_*p*_, *V*) diagrams shown in Figs 7d, e, f respectively (see **Supplementary Information** for details). In Fig. 7d, beginning at the magenta circle, the (*h*_*p*_,*V*) trajectory (magenta trace) moves to the right as *h*_*p*_ recovers during an IBI, until a burst onset threshold is crossed; point where the blue circles meet the red and black curves (see **Supplementary Information** for details), and eventually bursting begins; see upward arrow marking a jump-up in V at the onset of burst. During a burst, while V jumps up and-down during spikes, *h*_*p*_ moves to the left as slow inactivation accumulates during bursting (left arrow). Finally, when *h*_*p*_ reduces sufficiently, (*h*_*p*_, *V*) gets closer to the burst offset threshold (points at which the green and blue circles meet), and the burst terminates (down arrow). What is key in this figure is that the IBI is well-defined as the time period in which the (*h*_*p*_,*V*) trajectory moves along the red curve of steady states during the recovery process and moves past the burst onset threshold until a burst begins. However, when stochastic influences are present, the recovery period near-threshold is subject to random perturbations in V and can cause abrupt jump-up/spikes during the recovery period (see expanded inset in Fig. 7b). Projecting (*h*_*p*_,*V*) during this period on to Fig 7e, we note that the near-threshold noise amplitudes can occasionally push the (*h*_*p*_, *V*) trajectory (magenta) above a green region of attraction and this results in such abrupt spikes. Now, when *g*_*NaR*_ is added, the apparent restoration of burst regularity (see Fig. 7c) can be attributed to an expansion in this green shaded region as shown in Fig. 7f (see arrow pointing to a noise-tolerant region). In this situation, near-threshold random perturbations have less of an effect during the recovery process to induce abrupt spikes. This way, the net effect of *I*_*NaR*_ on slow Na^+^ inactivation prevents abrupt transitions into spiking regime following burst offset and in turn contributes to burst refractoriness and noise tolerance. We suggest that such a mechanism can make random fluctuations in membrane potential less effective in altering the precision of bursts and therefore aid information processing.

**Figure 7.**
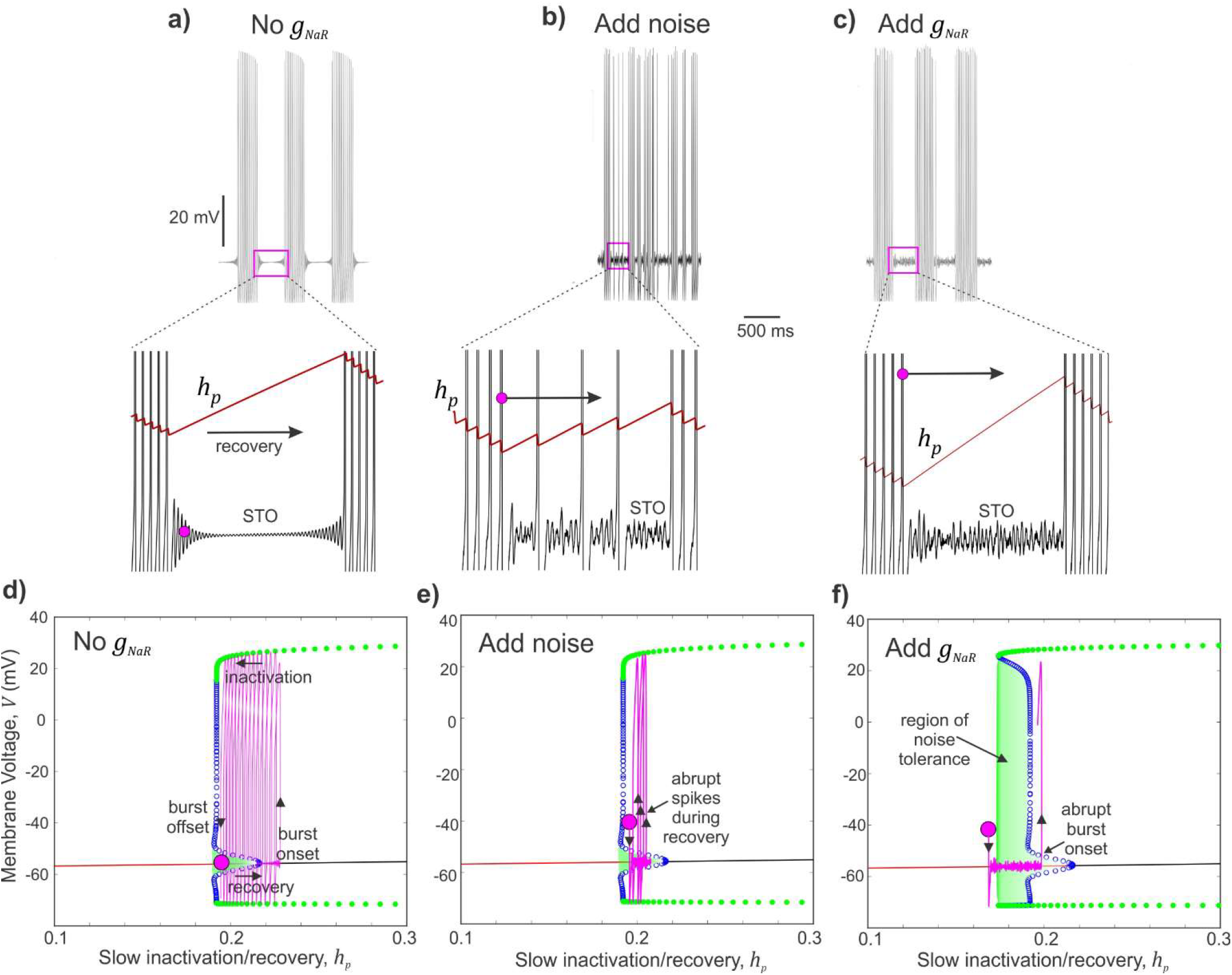
Resurgent Na^+^ current offers burst stability and noise modulation. **a - c)** Simulated membrane voltage shows neural activity patterns without any *g*_*NaR*_ **(a)**, with additive Gaussian noise input **(b)**, and with subsequent addition of *g*_*NaR*_ **(c)**. Expanded regions in **(a-c)** show membrane voltage sub-threshold oscillations (STO) during an inter-burst interval. Also overlaid is the evolution of the slow sodium inactivation/recovery variable; arrow indicates recovery during IBI, and magenta circle marks an arbitrary time point used to track these trajectories in **(d - f). d - f)** Bifurcation diagrams with projected trajectories of (*h*_∞_,*V*), shown in magenta to highlight the effect of addition of noise near sub-threshold voltages **(e)** and then the enlarged region of noise tolerance (highlighted green shaded region) due to addition of *g*_*NaR*_ in **(f)**. The magenta circle marks the beginning time points of each trajectory.

### Resurgent Na^+^ moderates burst entropy

The spike/burst intervals, their timing precision and order are important for information coding^38–42^. Given our prediction that *I*_*NaR*_ can offer noise tolerance and stabilize burst discharge, we examined whether it can reduce uncertainty in spike/burst intervals and restore order in burst discharge. We tested this using model simulations and dynamic-clamp experiments as shown in Fig. 8a- d. In both cases, as shown by spike raster plots in Figs. 8e and f, we disrupted the inter-event intervals (IEIs) by additive White/Weiner noise input while driving rhythmic burst discharge using step depolarization (also see **Methods**). Subsequent addition of *g*_*NaR*_ conductance restored the regularity of rhythmic bursting. We used Shannon’s entropy as a measure of uncertainty in IEIs and show that increases in entropy due to noise addition was reduced to control levels by subsequent increases in *g*_*NaR*_ as shown in Fig. 8g (see **Methods**). We also quantified the Coefficient of Variation (CV) and noted that adding noise which shortened IBIs, indeed decreased the CV, due to a reduction the standard deviation (s.d.) of the IEI distribution. Subsequent addition of *I*_*NaR*_, which significantly lengthened the IBIs, resulted in increases in CV values due to an increase in IEI s.d. Taken together, *I*_*NaR*_ moderated burst entropy and improved the regularity of spike/burst intervals.

**Figure 8.**
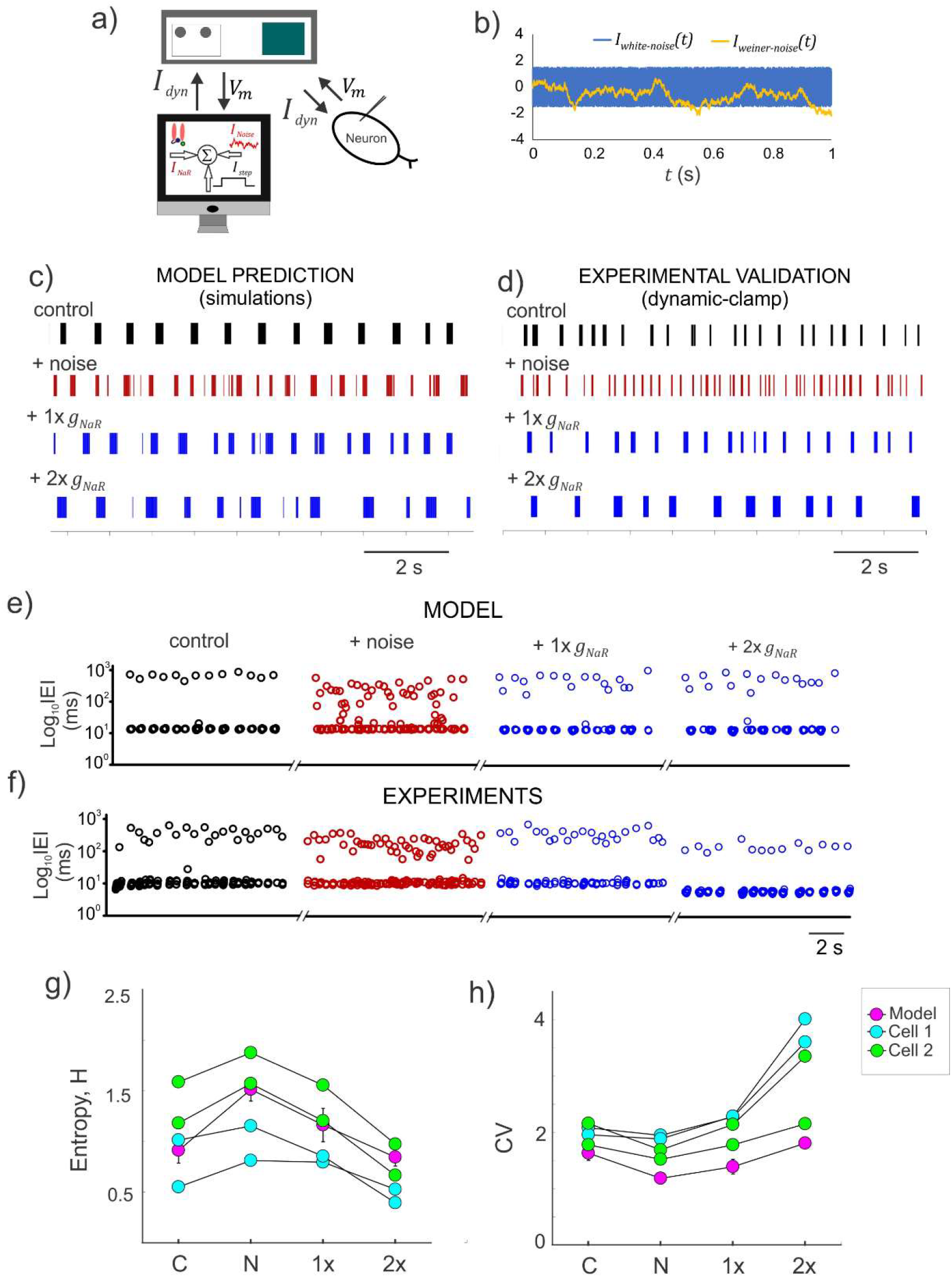
Resurgent Na^+^ current moderates the entropy of burst discharge. **a)** Schematic showing the experimental setup for real-time application of *I*_*NaR*_ and a stochastic input, *I*_*Noise*_ (see **Methods**); the dynamic-clamp current, *I*_*dyn*_ is the instantaneous sum of *I*_*NaR*_, a step current (*I*_*step*_) and *I*_*Noise*_. **b)** Simulated time series of two stochastic noise profiles used to disrupt the rhythmic burst discharge in Mes V neurons; *I*_*White-nois*_ was generated from a uniformly distributed random number and *I*_*weiner-noise*_ was generated from a normally distributed random number (see **Methods**). **c - d)** Raster plots showing patterns of inter-event intervals (IEIs) for the different conditions shown in the model **(e)**, and during dynamic-clamp **(f). e - f)** Time series of IEI shown on a log scale for the different conditions shown in the model **(e)**, and during dynamic-clamp **(f). g - h)** Shannon entropy (H) and coefficient of variability (CV) measured for IEIs under the different conditions presented in **(c)** and **(d)**. Plotted circles for the model represent an average across 10 trials, while individual trials are presented for the data points from two cells. In both **(g)** and **(h)**, C: control, N: after addition of random noise, 1x and 2x are supplements in *g*_*NaR*_ values.

## Discussion

Using a combination of mathematical modeling and simulations, theory and dynamic-clamp electrophysiology, we demonstrate a novel role for voltage-gated Na^+^ currents in burst control, noise modulation and information processing in sensory neurons. While the subclasses of Na^+^ currents presented here are experimentally inseparable, our phenomenological simplification using the HH-formalism with *in silico knock-in* of each Na^+^ current component using dynamic-clamp experiments, allowed examination of their individual contributions to burst control. Additionally, theoretical analysis suggested a putative negative feedback loop between the persistent inactivation and resurgent unblocking processes of the Nav1.6 Na^+^ channels. These results portend the apparent consequences on burst control and signal processing capacity of neurons when these currents are present.

### Stabilization of burst discharge by *I*_*NaR*_

In contrast with *I*_*NaP*_, which drives near threshold behavior and burst generation, *I*_*NaR*_ facilitated slow channel inactivation as bursts terminate (also note^11^). Increased channel inactivation due to *I*_*NaP*_ in turn prolonged the recovery from inactivation required to initiate subsequent burst of activity. Such an interaction between open-channel unblock process underlying *I*_*NaR*_, and, the slow inactivation underlying *I*_*NaP*_, offer a closed-loop *push-pull* modulation of ISIs and IBIs. Specifically, presence of *I*_*NaR*_ facilitates slow Na^+^ inactivation as shown by our theoretical analyses of model behavior; such enhanced slow channel inactivation can eventually shut off channel opening and unblocking. This resulted in burst stabilization. Theoretically, this represented an enlarged separatrix (or boundary) for transitioning from a sub-threshold non-spiking behavior to spiking behavior (see enlarged green shaded region in Fig. 7f), and the neuron becomes refractory to burst generation and hence offers noise tolerance.

Is this apparent effect of *I*_*NaR*_ physiologically plausible? Biophysical studies indicate that recovery from fast inactivation is facilitated in sodium channels that can pass resurgent current^30^; as shown here, this appears to be true for recovery from slow inactivation as well. Consistently, in the SCN8a knockout Med mouse, which lack the Nav 1.6 sodium channel subunit, recordings from mutant cells showed an absence of maintained firing during current injections, limited recovery of sodium channels from inactivation, and failure to accumulate in inactivated states. This is attributed to a significant deficit in *I*_*NaR*_ ^11, 20, 37^. Furthermore, maintained or repeated depolarization can allow a fraction of sodium channels in many neurons to enter inactivation states from which recovery is much slower than for normal fast inactivation (reviewed in^43^). Here, our simulations and model analyses predict that the presence, and increase in *I*_*NaR*_ conductance, provides for a such a physiological mechanism to maintain sustained depolarization and promote fast and slow Na^+^ inactivation.

### Sodium currents and signal processing in neurons

Neuronal voltage-gated Na^+^ currents are essential for action potential generation and propagation^5^. However, to enable fight-or-flight responses, an overt spike generation mechanism must be combined with *noise modulation* to extract behaviorally relevant inputs from an uncertain input space. Here we show that, the voltage-gated Na^+^ currents can serve an important role in neural signal processing (see summary in Fig. 9). As shown in the figure, a sub-threshold activated persistent Na^+^ current contributes to membrane resonance, a mechanism of bandpass filtering of preferred input frequencies^9^. We call this type of input gating, which is widely known to be important for brain rhythms^9, 41^, a *tune-in* mechanism (see figure legend). In some cases ambient noise or synaptic activity can amplify weak inputs and promote burst generation^44, 45^. This way, a tune-in mechanism such as the persistent Na^+^ current can contribute to weak input detection and promote burst coding^46, 47^ Then again, during rhythmic bursting, presence of resurgent Na^+^ maintains the order and precision of the timing events of bursts while preventing abrupt transitions into spiking phase due to stochastic influences as shown here. During ongoing sensory processing, such burst timing regulation can provide for noise cancellation or what we call a *tune-out* mechanism, which can mitigate random irregularities encoded in bursts (see Fig. 9 and **legend**). Whether this leads to improved sensory processing in the presence of natural stimuli and/or sensorimotor integration during normal behaviors needs to be validated. Our biological prediction here that a sensory neuron can utilize these voltage-gated Na^+^ currents as a *tune-in-tune-out* mechanism to gate preferred inputs, attenuate random membrane fluctuations and prevent abrupt transitions into spiking activity supports such a putative role.

**Figure 9.**
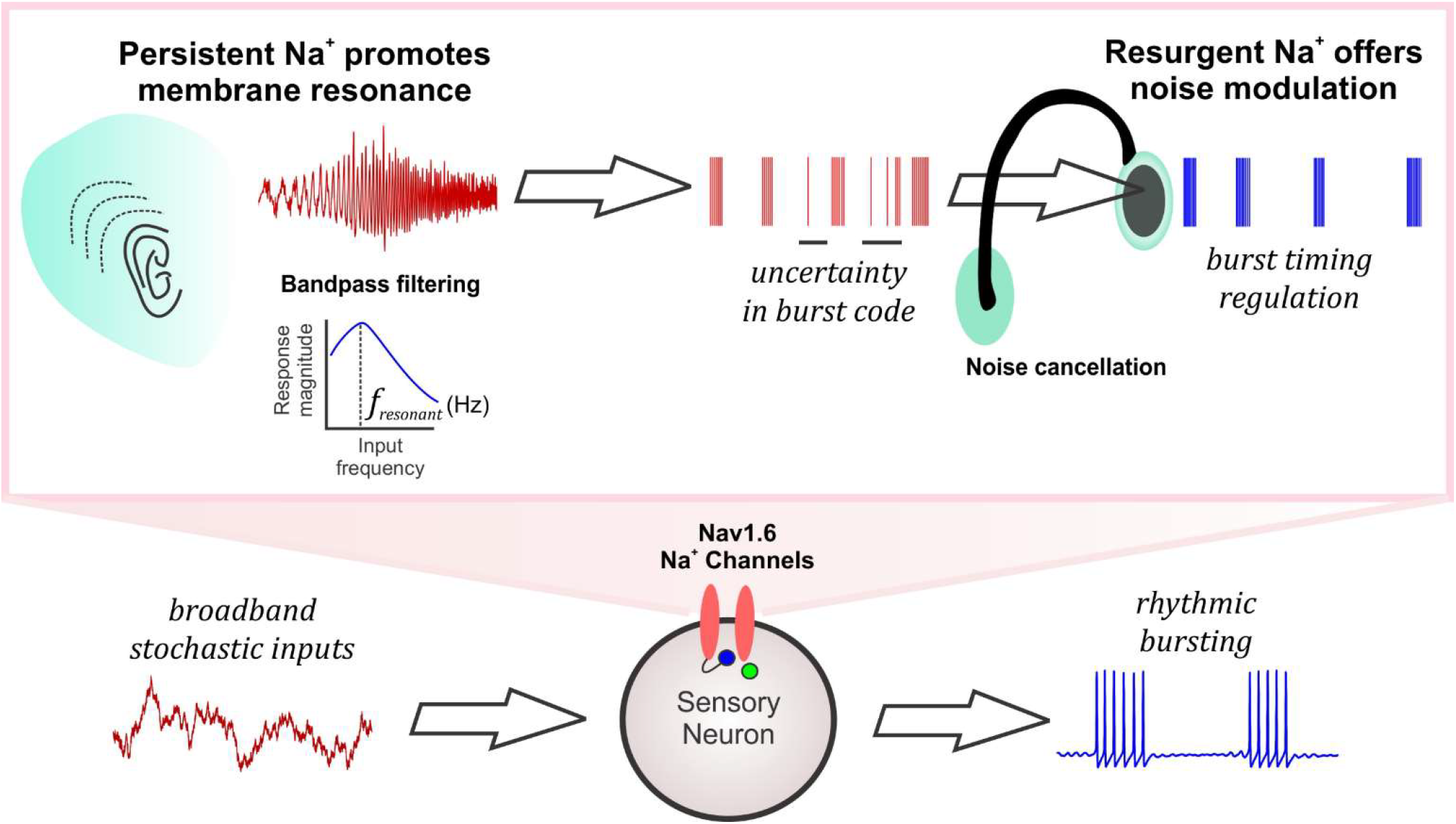
A consolidated role for persistent and resurgent Na^+^ currents in information processing. The schematic shows the broadband input space of a sensory neuron which can produce rhythmic bursting activity. The Nav1.6-type Na^+^ channels mediating the persistent Na^+^ current (schematized as a listening ear) enhance neuronal resonance and tune in relevant input frequencies by bandpass filtering (*f*_*resonance*_ refers to the resonant frequency of the neuron). Such selected inputs carrying behaviorally relevant stimuli are encoded as bursts but may be prone to uncertainty due to intrinsic or extrinsic stochastic influences. The resurgent Na^+^ current mediated by the Nav1.6 Na^+^ channels improve the regularity of spike/burst timing by mitigating the effects of noise by tuning out random spikes between bursts (schematized as a noise cancelling headphone), thus aiding in neural signal processing.

## Acknowledgements

We are grateful to Dr. Bruce Bean, Dr. Christopher Del Negro and Dr. Alan Garfinkel for helpful comments on an earlier version of the manuscript.

## Methods

### Neuron model for bursting activity

The conductance-based Mes V neuron model that we used to investigate the physiological role for *I*_*NaR*_ and *I*_*NaP*_ components of *I*_*Na*_ in burst discharge, incorporates a minimal set of ionic conductances essential for producing rhythmic bursting and for maintaining cellular excitability in these neurons ^8^. These include: 1) a potassium leak current, *I*_*leak*_, 2) sodium current, *I*_*Na*_ as described above, and, 3) a 4-AP sensitive delayed-rectifier type potassium current (*I*_*K*_) ^8,48^. The model equations follow a conductance-based Hodgkin-Huxley formalism ^5^ and are as follows.

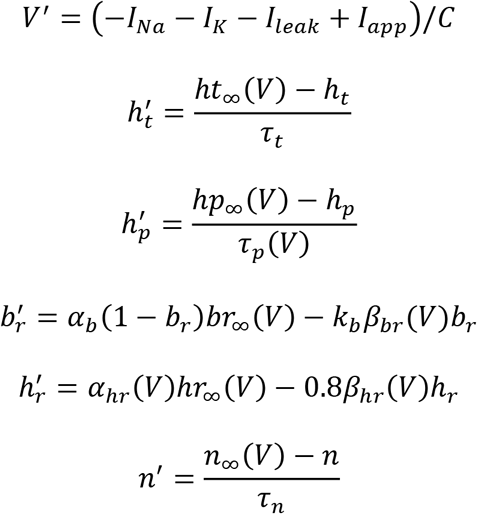

In what follows, we provide the formulation for each of the ionic currents and describe in detail, the novel *I*_*NaR*_ model.

#### A. Voltage-gated sodium currents

*In vitro* action potential clamp studies in normal mouse Mes V neurons, and voltage-clamp studies in Nav1.6 subunit SCN8a knockout mice have demonstrated existence of three functional forms of the total sodium current, *I*_*Na*_, including the transient (*I*_*NaT*_), persistent (*I*_*Nap*_) and resurgent (*I*_*NaR*_) components ^11, 14^. Each of these currents are critical for Mes V electrogenesis including burst discharge, however, their exclusive role is yet unclear. Lack of suitable experimental model or manipulation to isolate each of these TTX-sensitive components, led us to pursue an alternative approach involving computational model development of the physiological *I*_*Na*_. To further allow model-based experimental manipulation of individual components of the *I*_*Na*_, we designed a conductance-based model as follows. Although a single Nav1.6 channel can produce all three *I*_*Na*_ components observed experimentally, we used a set of three HH-type conductances, one for each of the transient, the persistent, and the resurgent components. This allowed us to easily manipulate these components independently to test their specific role in neural burst control. The equation for the total sodium current can be written as:

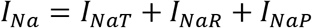

where,

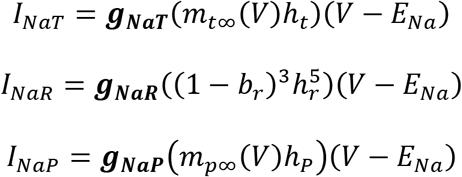

The maximal persistent conductance, *g*_*NaP*_ was set 5-10% of the transient, *g*_*NaT*_^49^ and the resurgent was set to 15-30% of *g*_*NaT*_, based on the relative percentage of maximum *I*_*NaR*_ and *I*_*NaT*_ as revealed by voltage-clamp experiments shown in Fig. 3; *E*_*Na*_ is the Na^+^ reversal potential.

Based on experimental data, the gating function/variable, *m*_*t∞*_(*V*), and *h*_*t*_, for *I*_*NaT*_, and, *m*_*p∞*_(*V*), and, *h*_*P*_, for *I*_*NaP*_ are modeled as described in ^8^. The rate equations for the inactivation gating variables *h*_*t*_, and, *h*_*P*_, model the fast and slow inactivation of the transient and persistent components respectively. The activation gates are steady-state voltage-dependent functions, consistent with fast voltage-dependent activation of *I*_*Na*_.

Steady-state voltage-dependent activation and inactivation functions of *transient* sodium current respectively include:

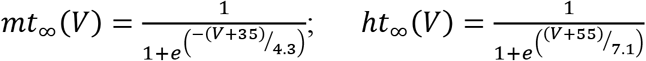

Steady-state activation, inactivation and steady-state voltage-dependent time constant of inactivation for *persistent* sodium current respectively include:

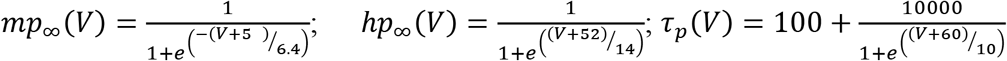

The novel *I*_*NaR*_ formulation encapsulates the block/unblock mechanism using a block/unblock variable (*b*_*r*_), and, a second hypothetical variable for a competing inactivation, which we call, *h*_*r*_. We call this a *hybrid* model, to highlight the fact that the model implicitly incorporates the history or state-dependent sodium resurgence, following a transient channel opening, and combines this into a traditional Hodgkin-Huxley type conductance-based formulation. In the 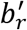 and 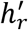 rate equations for *b*_*r*_, and, *h*_*r*_, the block/unblock variable, *b*_*r*_ increases or grows according to the term, *α*_*b*_(1 – *b*_*r*_)*br*_∞_(*V*), and decays as per the term, *k*_*b*_*β*_*br*_(*V*)*b*_*r*_, described as follows:

***α***_***b***_**(1 – *b*_*r*_)*br***_∞_**(*V*)**: In this growth term, we incorporate state-dependent increase in *b*_*r*_, as follows; we assume that the rate of increase in *b*_*r*_ is proportional to the probability of channels currently being in the open state, with a rate constant, *α*_*b*_ which we call ‘rate of unblocking’; such probability is a function of the membrane voltage given by, *br*_∞_(*V*), defined as below:

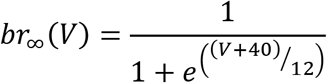

The term (1 – *br*_∞_(*V*)), models the steady-state voltage-dependency guiding the unblocking process. The channels being in open state is represented by the term, (1 – *b*_*r*_). Note that if (1 – *b*_*r*_) is close to 1, this means that larger proportion of channels are in an open state, and therefore *b*_*r*_ grows faster, promoting blocking. We modeled *br*_∞_(*V*) as a decreasing sigmoid function, such that, at negative membrane potentials, channels have a high probability to enter future depolarized states and therefore, (1 – *b*_*r*_) ~ 0, in turn, *b*_*r*_ does not grow fast.

***k***_**b**_***β***_***br***_(***V***)***b***_***r***_: In this decay term, we assume that the rate of decay of *b*_*r*_, is proportional to the probability of channels being in the blocked state, with a constant of proportionality *k*_*b*_, and, this probability is given by a voltage-dependent function, *β*_*br*_(*V*), defined as below:

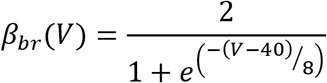

Note that, *β*_*br*_(*V*) gives a high probability at depolarized potentials, indicating a blocked state and enables decrease in *b*_*r*_ in subsequent time steps.

Taken together, *b*_*r*_, represents a phenomenological implementation of a previously described block/unblock mechanism of a cytoplasmic blocking particle^19^ (see schematic of channel gating in Fig. 3a). Additionally, a hypothetical competing inactivation variable, *h*_*r*_, sculpts the voltage-dependent rise and decay times and peak amplitude of sodium resurgence at −40 mV following a brief depolarization (i.e., transient activation), as observed in voltage-clamp experiments (see Fig. 3b). The functions, *α*_*hr*_(*V*),*β*_*hr*_(*V*) and *hr*_∞_(*V*) are defined as voltage-dependent rate equations that guide the voltage-dependent kinetics and activation/inactivation of the *I*_*NaR*_ component as given below.

The steady-state voltage-dependency of the competing inactivation necessary to generate a *resurgent* Na^+^ current is defined as follows:

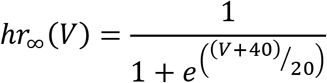

The voltage-dependent rate functions of such inactivation is defined by two functions as follows:

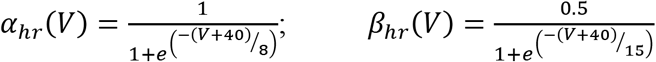

The steepness of the voltage-dependent sigmoid functions for activation and inactivation were tuned to obtain the experimentally observed *I*_*NaR*_ activation (see Fig. 3; also see ^11, 14, 30^). To obtain the kinetics (rise and decay times) of *I*_*NaR*_ comparable to those observed during voltage-clamp experiments (see **Supplementary Fig. 3**), the model required three units for the blocking variable ((1 – *b*_*r*_)^3^) and five units for the inactivation variable 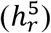 (see *I*_*NaR*_ equation). Together, the modeled *I*_*Na*_ reproduced the key contingencies of the Nav1.6 sodium currents (see **Supplementary Fig. 2**) ^18, 30, 50^.

Sensitivity analyses was conducted for the key parameters of *I*_*NaR*_ gating including *α*_*b*_, and, *k*_*b*_. Note that these two parameters control the rate of blocking. As expected, increasing *α*_*b*_, that controls rate of increase in *b*_*r*_, decreased the peak amplitude of *I*_*NaR*_, similar to an experimental increase in block efficacy by a *β*-peptide (e.g., ^19^). On the other hand, *k*_*b*_ also moderates *b*_*r*_, and increasing *k*_*b*_, enhances *b*_*r*_ decay rate, that significantly enhanced *I*_*NaR*_, and, therefore burst duration (not shown). Large increases in *k*_*b*_ significantly enhanced *I*_*NaR*_, and indeed transformed bursting to high frequency tonic spiking. However, the effects of *I*_*NaR*_ on bursting described in the results section were robust for a wide range of values of these parameters (>100% increase from default values), and, for our simulations, the range of values, *α*_*b*_ = 0.08 *to* 0.1, *k*_*b*_ = 0.8 *to* 1.2, were used to reproduce Mes V neuron discharge properties. To reproduce experimentally observed spike width, we additionally tuned the inactivation time constant, *τ*_*t*_ = 1.5 ± 0.5, for *I*_*NaT*_.

#### B. Potassium and leak currents

The 4-AP sensitive delayed-rectifier type potassium current, *I*_*K*_, and the leak current, *I*_*leak*_ were modeled similar to ^8^ as below; also see ^48^.

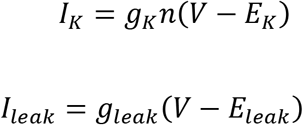

where, the steady-state voltage-dependent activation function for the gating variable, *n* is given as:

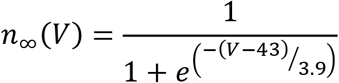

*E*_*k*_ and *E*_*leak*_ are K^+^ and leak reversal potentials respectively. Model parameter values used are provided in **Supplementary Table. 1**.

### Brain slice preparation

All animal experiments were performed in accordance to the institutional guidelines and regulations using protocols approved by Animal Research Committee at UCLA. Experiments were performed in P8-P14 wild-type mice of either sex. Mice were anesthetized by inhalation of isofluorane and then decapitated. The brainstem was extracted and immersed in ice-cold cutting solution. The brain-cutting solution used during slice preparation was composed of the following (in mM): 194 Sucrose, 30 NaCl, 4.5 KCl, 1.2 NaH_2_PO_4_, 26 NaHCO_3_, 10 glucose, 1 MgCl_2_. The extracted brain block was mounted on a vibrating slicer (DSK Microslicer, Ted Pella) supported by an agar block. Coronal brainstem sections consisting of rostro-caudal extent of Mes V nucleus, spanning midbrain and pons were obtained for subsequent electrophysiological recording.

### Voltage-clamp electrophysiology

To obtain direct experimental data to drive *I*_*NaR*_ model development, we performed voltage-clamp experiments on Mes V neurons and recorded Na^+^ currents by blocking voltage-gated K^+^ and Ca^2+^ currents similar to ^11^. The pipette internal solution contained the following composition (in mM): 130 CsF, 9 NaCl, 10 HEPES, 10 EGTA, 1 MgCl_2_, 3 K_2_-ATP, and 1 Na-GTP. The external recording solution contained the following composition (in mM): 131 NaCl, 10 HEPES, 3 KCl, 10 glucose, 2 CaCl_2_, 2 MgCl_2_, 10 tetraethylammonium (TEA)-Cl, 10 CsCl, 1 4-aminopyridine (4-AP), and 0.3 CdCl_2_. The voltage-clamp protocol consisted of a holding potential of −90 mV followed by a brief voltage pulse (3 ms) of +30 mV, to remove voltage-dependent block, followed by voltage steps between −70 mV to −10 mV, in steps of 10 mV for ~ 100 ms to activate *I*_*NaR*_, and then returned to −90 mV. A 1 μM TTX abolished the Na^+^ current and the residual leak current was subtracted to isolate evident sodium currents. Recordings with series resistance *R*_*series*_ > 0.1*R*_*m*_ were discarded, where *R*_*m*_ is the input resistance of the neuron; we did not apply any series resistance compensation.

### Dynamic-clamp electrophysiology

Dynamic-clamp electrophysiology and *in vitro* current-clamp recording were used for testing the physiological effects of Na^+^ currents on burst discharge as well as noise-mediated entropy changes corrected by *I*_*NaR*_ ^51^. We selected neurons responding with a bursting pattern in response to supra-threshold step current injection in the Mes V nucleus in brainstem slice preparation for our study; >50% of neurons showing other patterns (e.g., tonic or single spiking cells) were discarded. Dynamic-clamp was successfully performed in bursting cells (n = 10). For dynamic-clamp recording, slices were placed in normal ACSF at room temperature (22-25°C). The ACSF recording solution during patch-clamp recording consisted of the following (in mM): 124 NaCl, 4.5 KCl, 1.2 NaH_2_PO_4_, 26 NaHCO_3_, 10 glucose, 2 CaCl_2_, 1 MgCl_2_. Cutting and recording solutions were bubbled with carbogen (95% O_2_, 5% CO_2_) and maintained at pH between 7.25 - 7.3. The pipette internal solution used in current clamp experiments was composed of the following (in mM): 135 K-gluconate, 5 KCl, 0.5 CaCl_2_, 5 HEPES (base), 5 EGTA, 2 Mg-ATP, and 0.3 Na-ATP with a pH between 7.28 - 7.3, and osmolarity between 290 ± 5 mOsm. Patch pipettes (3 - 5 MΩ) were pulled using a Brown/Flaming P-97 micro pipette puller (Sutter Instruments). Slices were perfused with oxygenated recording solution (~2ml/min) at room temperature while secured in a glass bottom recording chamber mounted on an inverted microscope with differential interface contrast optics (Zeiss Axiovert 10). Current clamp (and dynamic-clamp) data were acquired and analyzed using custom-made software (G-Patch, Analysis) with sampling frequency: 10 kHz; cut-off filter frequency: 2 kHz.

The Linux-based Real-Time eXperimental Interface (RTXI v1.3) was used to implement dynamic-clamp, running on a modified Linux kernel extended with the Real-Time Applications Interface, which allows high-frequency, periodic, real-time calculations ^52^. The RTXI computer interfaced with the electrophysiological amplifier (Axon Instruments Axopatch 200A, in current-clamp mode) and the data acquisition PC, via a National Instruments PCIe-6251 board. Euler’s method with step size 0.05 ms was used for model integration resulting in a computation frequency of 20 kHz.

The model *I*_*NaR*_ current used for dynamic clamping into Mes V neuron *in vitro* was developed as discussed above. The ionic conductance *g*_*NaR*_ was set to suitable values to introduce model *I*_*NaR*_ current into a Mes V neuron during whole-cell current-clamp recording. For dynamic-clamp experiments involving *g*_*NaR*_ mediated noise modulation, two approaches were used to model random noise *I*_*noise*_, generated in RTXI and injected as pA current:

1. Uniform-distributed random values (white noise):

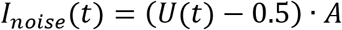

where *U*(*t*) was a uniformly-distributed random number between 0 and 1 generated by the C^++^ *rand()* function, and *A* was a scaling factor, in this case determining peak-to-peak noise amplitude. It was varied between 3 and 5, to adjust noise amplitudes to produce discernable burst irregularities. The mean injected noise was 0 pA, and its effective standard deviation varied between 0.87 pA (A=3) and 1.44 pA (A=5).
2. Normally distributed random values (Wiener noise)

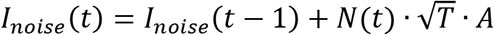

where *t* indicates the value generated at the current computation cycle, and *t* − 1 to the value of the previous computation cycle. *T* was the computation period in seconds (0.00005 s for 20 kHz computation) and was discretized to mimic a Weiner process (random-walk); *A* was a scaling factor, in this case affecting the noise increment each computation cycle, varying between 3 and 5. *N(t)* was a normally-distributed random number with mean, *μ* = 0, and standard deviation, *σ* = 1. *N(t)* was generated from C^++^-generated uniformly-distributed random numbers using the central limiting theorem:

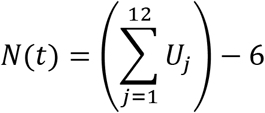

where each *U*_*j*_ is a uniformly-distributed random number between 0 and 1 generated using the C^++^ *rand()* function. In three 1-second simulations, this noise had mean −0.31±0.19 pA (*A*=3), −0.21±0.62 pA (*A*=4) or −0.59±0.33 pA (*A*=5); and standard deviation 0.77±0.13 pA (*A*=3), 1.07±0.15 pA (*A*=4) or 1.68±0.44 pA (*A*=5). We used stochastic current injection as an external input (additive noise) in order to produce irregularities/uncertainties in burst discharge. Our choice of the noise model was to experimentally disrupt spike timing regularity^45^ and is not directly based on any known noise characteristics in Mes V neurons. The jaw muscle spindle afferent Mes V neurons are not known to have spontaneous synaptic events and we did not characterize Na^+^ channel fluctuations in these neurons^53^. Nonetheless, the stochastic noise we used as shown in the representative example in Fig. 8 most closely matched a diffusive synaptic noise model with Gaussian distribution^54^; Fig. 8c illustrates the temporal features of the noise inputs described above. A Gaussian white noise generated in MATLAB with zero mean and unit standard deviation or a Weiner noise generated in XPPAUT were used to disrupt firing patterns in the model neuron.

### Model simulation and Data analyses

Model simulation and all the analyses were performed using MATLAB (Mathworks™) (see model code provided as **Supplementary Information**). Model bifurcation analyses were performed using XPPAUT/AUTO ^55^. A variable step Runge-Kutta method ‘ode45’ was used for current-clamp simulations and ‘ode23s’ was used for voltage-clamp simulations.

Inter-event intervals (IEI) between spikes in dynamic-clamp recordings were detected using Clampfit 9.0 software and were classified *post hoc* as ISIs and IBIs based on a bi-modal distribution of IEIs. Typically, IEI values < 40 ms were considered as ISIs within bursts and IEI values ≥ 40 ms were considered as IBIs. Any occasional isolated spikes were eliminated from analyses for burst duration calculations.

To calculate Shannons’ entropy^56^ in the inter-event intervals (IEIs), we generated histograms of and calculated the probabilities for each bin of the underlying IEI distributions for each 10 sec spike trains. The probability of *k*^*th*^ IEI bin from a distribution of *n* equal size bins was calculated from the bin counts, *N(k)* as:

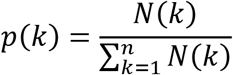

The entropy, H was calculated using the following formula:

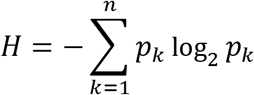

where, *n* is the total number of IEI bins, each with probability, *p*_*k*_.

The coefficient of variation (CV) in IEIs was calculated as follows:

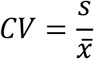

where, *s* is the IEI sample standard deviation, and, 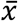 is the sample mean.

## Supplementary Information

**Supplementary Figure 1. a)** A state-based Markov model for *I*_Na_ as in Raman & Bean (2001). C1 to C5 represent sequential closed states and O denotes the open state. I1 through I6 represent the inactivation states for the normal inactivation mechanism. The rate constants in ms^−1^ are: *α* = 190*e*^(*V/20*)^, *β* = 1.6*e*^(*-V*/19.5)^, *γ* = 190, and, *δ* = 41. The normal inactivation is voltage-independent and occurs with rate constants, C_on_ = 0.002 ms^−1^, C_off_ =0.53 ms^−1^, while the O_on_ and O_off_ are 0.2 and 0.003 ms^−1^ respectively. The binding factor ‘*a*’ for the inactivating particle is given by (C_off_/C_on_)/(O_off_/O_on_))^1/8^. The rate constant *ε* = 1.75 ms^−1^, is the rate at which the open channel block occurs and the exit from the blocked state is modeled as a voltage-dependent process given by ζ = 0.03*e*^4/67^ ms^−1^. b) Voltage-clamp simulation of the Markov model shows the resurgent sodium current; Protocol is shown in the inset; Also highlighted are the transient and persistent components. c) The sodium current generated during action potentials both in our three-component Hodgkin-Huxley (3HH) type model and the Markov model are shown for comparison; spikes are in blue and sodium current is in red.

**Supplementary Figure 2. a)** Model simulation of sodium channel recovery and its relationship with resurgent current flow. Upper traces in (a) and (b) show two voltage-clamp protocols with voltages as shown; Δ*t* represents variable intervals from 3 – 39 ms for the test voltage of −40 mV; lower traces show the corresponding current responses. c) Percent channel availability calculated as the peak current at the test steps to −40 mV, normalized to the peak current at the reference pulse of 0 mV; d) Steady-state availability, calculated as peak current evoked at 0 mV following a 200 ms pre-pulse conditioning steps (−90 to +30 mV), normalized by the peak current at 0 mV following conditioning at −90 mV.

**Supplementary Figure 3. a)** Rise and decay kinetics in the model (magenta) and experimental trace (black) at a test potential of −40 mV where maximum resurgent current is seen. b, c) Rise and decay times for the model and experiments (n=5) are shown for three test potentials at which a sizable current was seen in ≥ 50% of the time; Rise time was calculated from 10 – 90% of the peak value, and, decay time was calculated from 90 – 10% of steady-state value at the end of the test pulse. Error bars indicate standard deviation.

**Supplementary Figure 4. a, c)** Representative burst discharge showing the effects of dynamic-clamp subtraction of *g*_NaR_ (a) and *g*_NaP_ (c); decreasing values of 0.25 and 0.5 nS/pF, and 0.05 and 0.1 nS/pF were used for 1X and 2X *g*_NaR_ and *g*_NaP_ respectively. Red double arrows highlight inter-burst intervals (IBIs). The magenta rectangles and the corresponding numbers are shown in expanded time and voltage in the boxed inset to highlight abolition of STO due to *g*_NaP_ subtraction (compare upper and middle traces in the box), which was further comparable to setting *g*_NaP_ = 0 in neuron model simulations (lower trace). a, c) Box plots showing effects of *g*_NaR_ (b) and *g*_NaP_ (d) subtraction on inter-spike intervals (ISIs) within bursts; C is control. A one-way ANOVA for treatment effect on ISIs had a *p* < 0.001 for *g*_NaR_ subtraction, with no significant effect of *g*_NaP_ between C and −1x; however, all spikes were abolished upon −2x *g*_NaP_ application. Asterisks in (b) indicate statistical significance with *p* < 0.001 using a Student t-test for group comparisons (Control v/s −1x and −1x v/s −2x).

## Supplementary Text

### Bifurcation Analyses of Model Behavior

Mathematical analytical approaches enable examination of experimentally intractable mechanisms and provide deeper insights into the behavior of complex nonlinear dynamical systems. We combined model simulations with geometric dynamical systems methods to examine the *I*_*NaR*_’s mechanism of burst control. In this supplementary text, we explain the theoretical concepts underlying the so-called bifurcation diagrams presented in Figs. 6d and e. First, to generate theses diagrams, we dissected the system of equations representing the bursting neuron model into fast and slow subsystems^2^. This enabled understanding the qualitative changes in the behavior of fast subsystem consisting of the membrane potential (*V*) due to changes in the slow kinetics of persistent Na^+^ inactivation/recovery (*h*_*p*_) (see Figs. 6a-c highlighting relative timescales of *V* and *h*_*p*_ during bursting activity). Note that during a burst of activity, *h*_*p*_ is inactivating (decreasing) during each spike, however, recovers from inactivation on a relatively slow timescale. In the diagrams in Figs. 6d and e, we treat the slow variable *h*_*p*_ as a parameter that when slowly changes, causes qualitative changes in the fast variable, *V*. For example, we can determine for which values of *h*_*p*_ the fast subsystem generates either near rest behavior or quiescence (stable equilibria) or spikes (stable oscillations) or both. Note that there are two values of *h*_*p*_, *h*_HB_ (onset threshold) and *h*_SNP_ (offset threshold), such that *V* exhibits: i) a near rest behavior (stable equilibrium – red curve) for *h*_*p*_ < *h*_HB_; ii) an unstable equilibrium for *h*_*p*_ > *h*_HB_; and iii) spiking for *h*_*p*_ < *h*_SNP_ (stable oscillations – green circles). As *h*_*p*_ gradually increases during IBI, indicating channel recovery, the membrane potential, *V* transitions from a resting state (or stable equilibria) until it reaches the Andronov-Hopf bifurcation point, labeled “onset threshold” (e.g., Figs. 6d), that is representative of spike onset threshold. Past this threshold, *V* becomes unstable and spiking phase begins whose amplitudes are bounded by the upper and lower green curves marking the maximum (peak) and minimum (trough) values of membrane potential, *V*. These represent stable oscillations (or periodic solution) when the neuron is in the bursting phase. For, *h*_SNP_ < *h*_*p*_ < *h*_HB_, the membrane potential, *V* shows bistability: for those values of *h*_*p*_, there is both a stable equilibrium and a stable oscillation. The same diagrams are reproduced for specific *g*_NaR_ and *g*_*NaP*_ values in Figs. 7d-f where the green shaded region that represents a region of attraction for the stable curve of equilibria are highlighted. Physiologically, this region houses the sub-threshold oscillations that decay in amplitude as a burst terminates.

