## Supplementary Material for "Resurgent Na^+^ Current Offers Noise Modulation in Bursting Neurons"

**Supplementary Information**

**4** Supplementary figures and legends

**2** Tables

**1** piece of Supplementary information providing the *Model analysis using dynamical systems methods*

**2** pieces of Supplementary information providing the *Model code as MATLAB scripts*

**3** pieces of Supplementary information providing the *Dynamic-clamp C^++^* *code*

**Supplementary Figure 1**

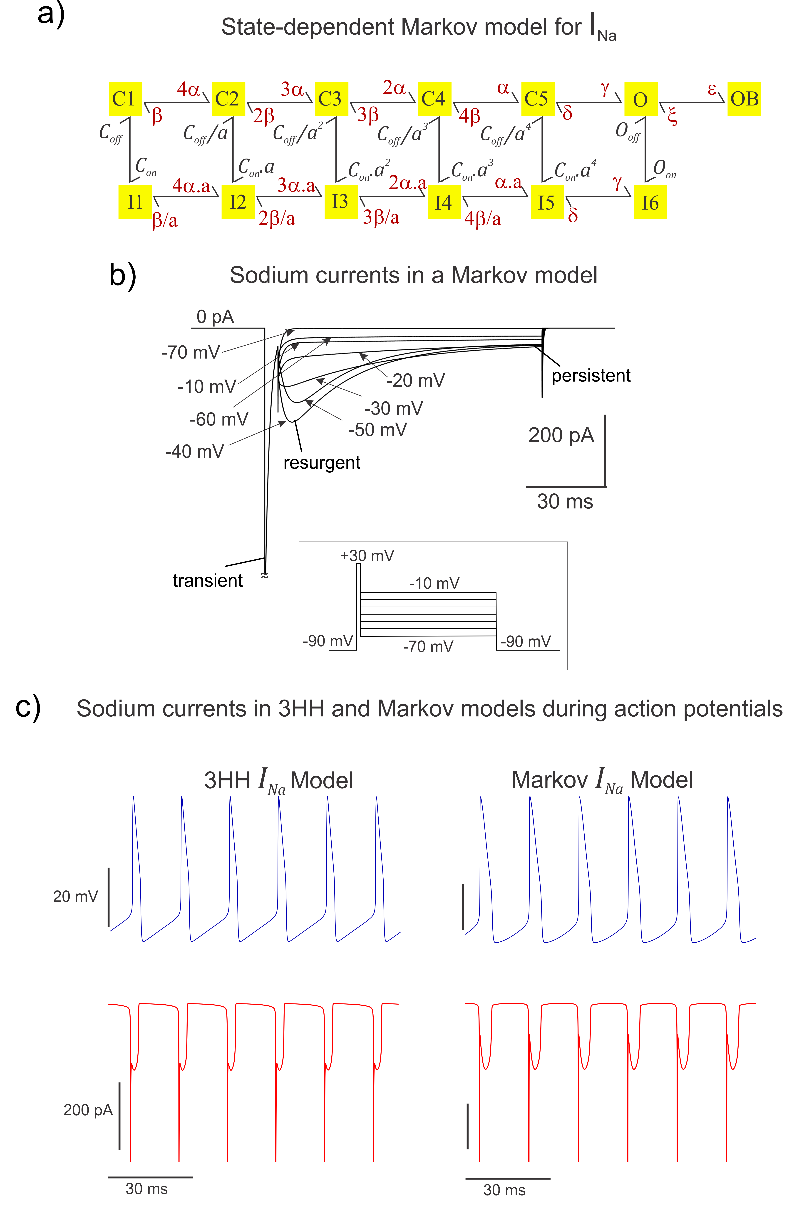

**a)** A state-based Markov model for $I_{Na}$ as in Raman & Bean (2001). C1 to C5 represent sequential closed states and O denotes the open state. I1 through I6 represent the inactivation states for the normal inactivation mechanism. The rate constants in ms^-1^ are: $\alpha=190e^{(V/20)}$, $\beta=1.6e^{(-V/19.5)}$, $\gamma=190$, and, $\delta= 41$. The normal inactivation is voltage-independent and occurs with rate constants, C_on_ = 0.002 ms^-1^, C_off_ =0.53 ms^-1^, while the O_on_ and O_off_ are 0.2 and 0.003 ms^-1^ respectively. The binding factor ‘$a$’ for the inactivating particle is given by (C_off_/C_on_)/(O_off_/O_on_))^1/8^. The rate constant 𝜀 = 1.75 ms^-1^, is the rate at which the open channel block occurs and the exit from the blocked state is modeled as a voltage-dependent process given by ζ = 0.03$e^{4/67}$ ms^-1^. **b)** Voltage-clamp simulation of the Markov model shows the resurgent sodium current; Protocol is shown in the inset; Also highlighted are the transient and persistent components. **c)** The sodium current generated during action potentials both in our three-component Hodgkin-Huxley (3HH) type model and the Markov model are shown for comparison; spikes are in blue and sodium current is in red.

**Supplementary Figure 2**

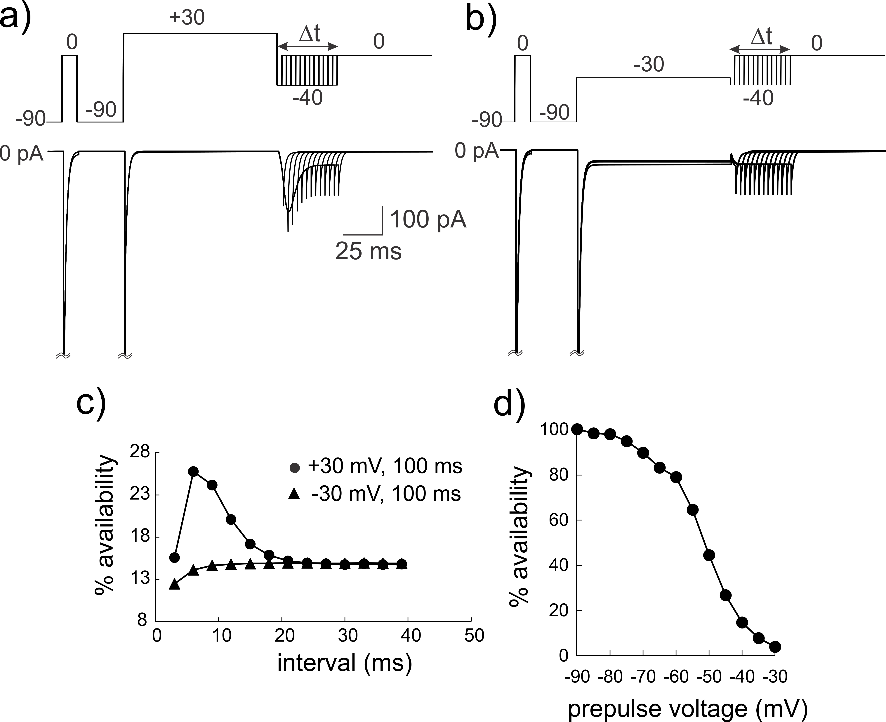

**a)** Model simulation of sodium channel recovery and its relationship with resurgent current flow. Upper traces in **(a)** and **(b)** show two voltage-clamp protocols with voltages as shown; Δ𝑡 represents variable intervals from 3 – 39 ms for the test voltage of -40 mV; lower traces show the corresponding current responses. **c)** Percent channel availability calculated as the peak current at the test steps to -40 mV, normalized to the peak current at the reference pulse of 0 mV; **d)** Steady-state availability, calculated as peak current evoked at 0 mV following a 200 ms pre-pulse conditioning steps (-90 to +30 mV), normalized by the peak current at 0 mV following conditioning at -90 mV.

**Supplementary Figure 3**

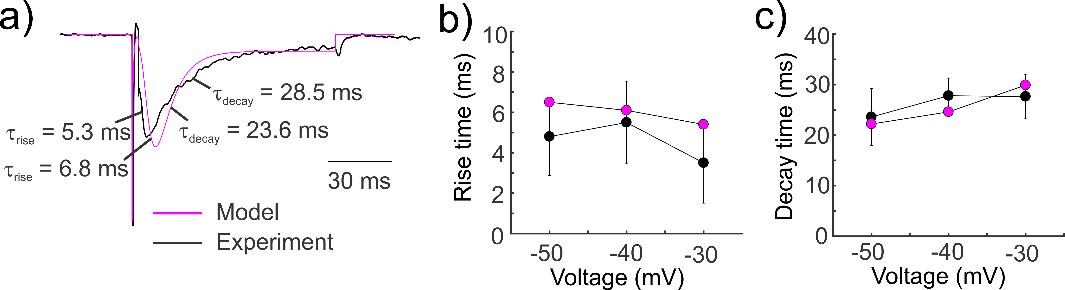

**a)** Rise and decay kinetics in the model (magenta) and experimental trace (black) at a test potential of -40 mV where maximum resurgent current is seen. **b, c)** Rise and decay times for the model and experiments (n=5) are shown for three test potentials at which a sizable current was seen in ≥ 50% of the time; Rise time was calculated from 10 – 90% of the peak value, and, decay time was calculated from 90 – 10% of steady-state value at the end of the test pulse. Error bars indicate standard deviation.

**Supplementary Figure 4**

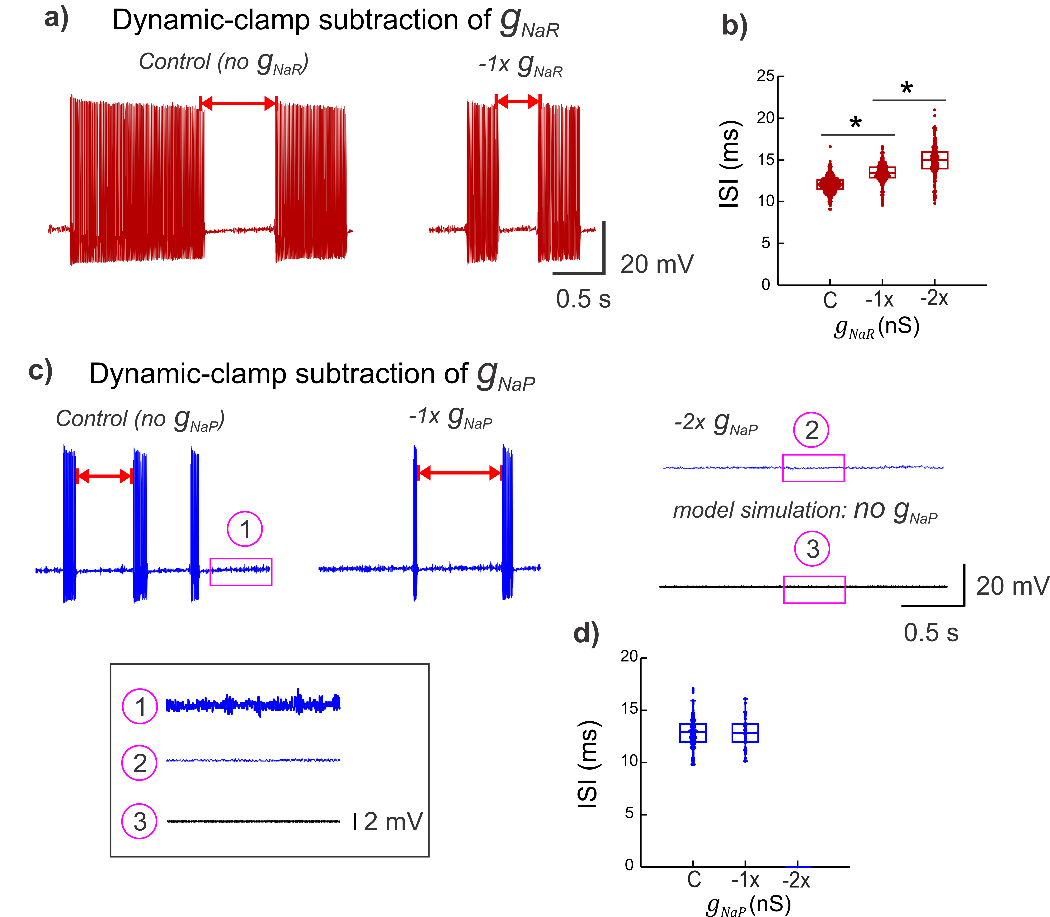

**a, c)** Representative burst discharge showing the effects of dynamic-clamp subtraction of $g_{NaR}$ **(a)** and $g_{NaP}$ **(c)**; decreasing values of 0.25 and 0.5 nS/pF, and 0.05 and 0.1 nS/pF were used for 1X and 2X $g_{NaR}$ and $g_{NaP}$ respectively. Red double arrows highlight inter-burst intervals (IBIs). The magenta rectangles and the corresponding numbers are shown in expanded time and voltage in the boxed inset to highlight abolition of STO due to $g_{NaP}$ subtraction (compare upper and middle traces in the box), which was further comparable to setting $g_{NaP}=0$ in neuron model simulations (lower trace). **a, c)** Box plots showing effects of $g_{NaR}$ **(b)** and $g_{NaP}$ **(d)** subtraction on inter-spike intervals (ISIs) within bursts; C is control. A one-way ANOVA for treatment effect on ISIs had a $p<0.001$ for $g_{NaR}$ subtraction, with no significant effect of $g_{NaP}$ between C and -1x; however, all spikes were abolished upon -2x $g_{NaP}$ application. Asterisks in **(b)** indicate statistical significance with $p<0.001$ using a Student t-test for group comparisons (Control v/s -1x and -1x v/s -2x).

**Supplementary Tables**

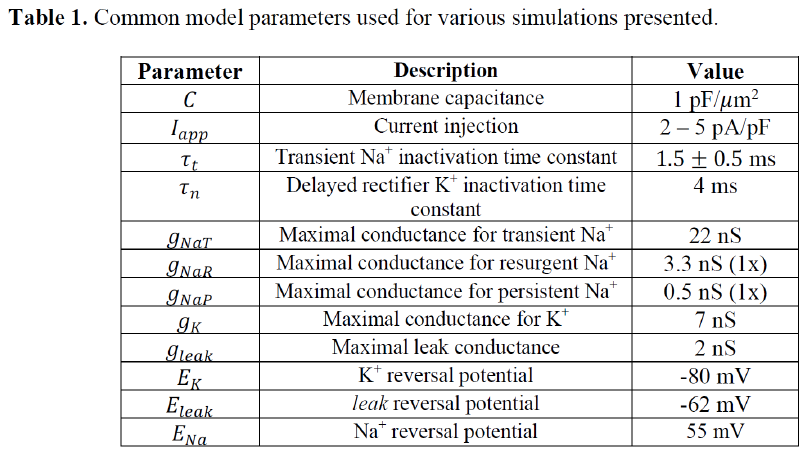

**Table 2.** Statistical summary of burst analysis for dynamic-clamp application of resurgent ($g_{NaR}$) and persistent ($g_{NaP}$) Na^+^ conductances/currents.

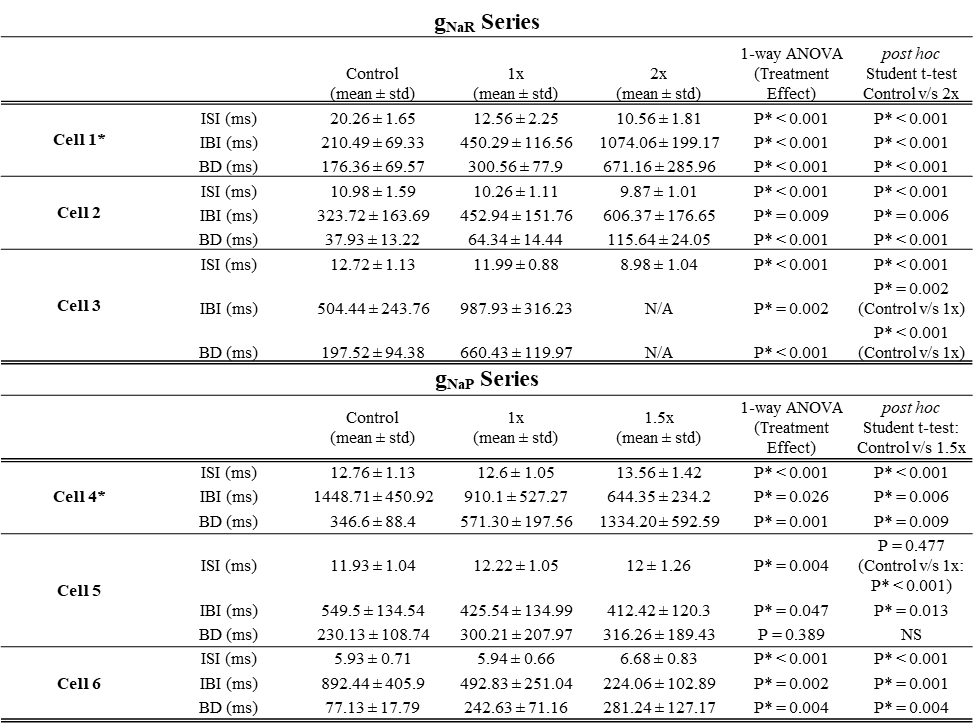

*Note*: $n \geq38$ for ISIs, $n \geq4$ for IBIs and BDs and varies for each cell and between each treatment.

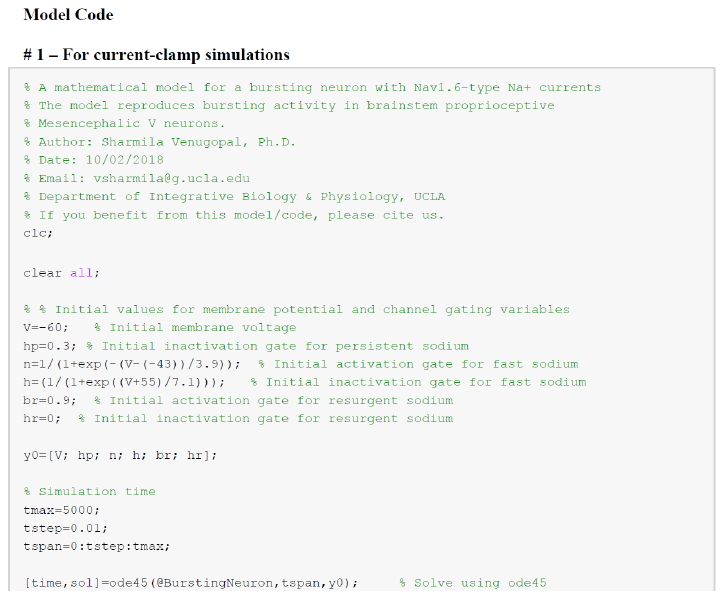

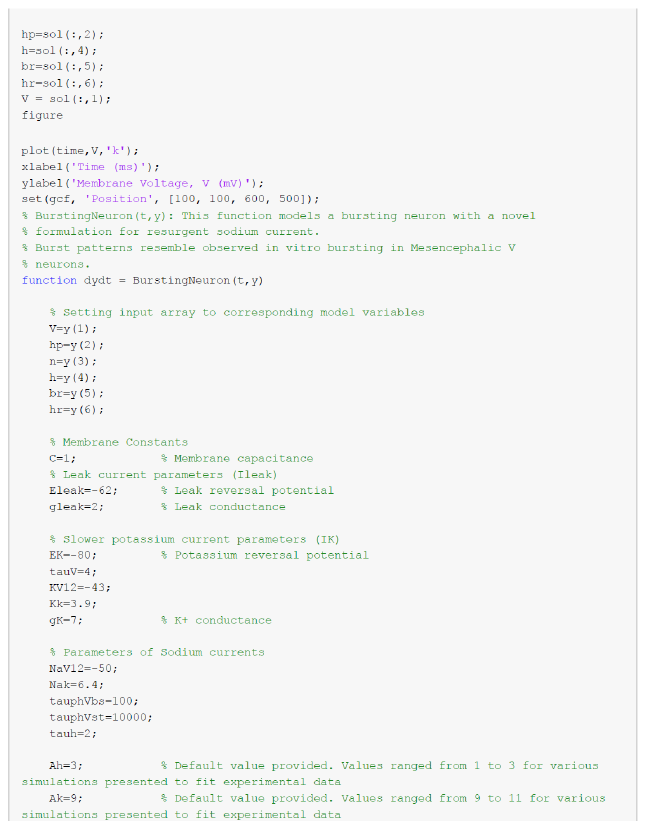

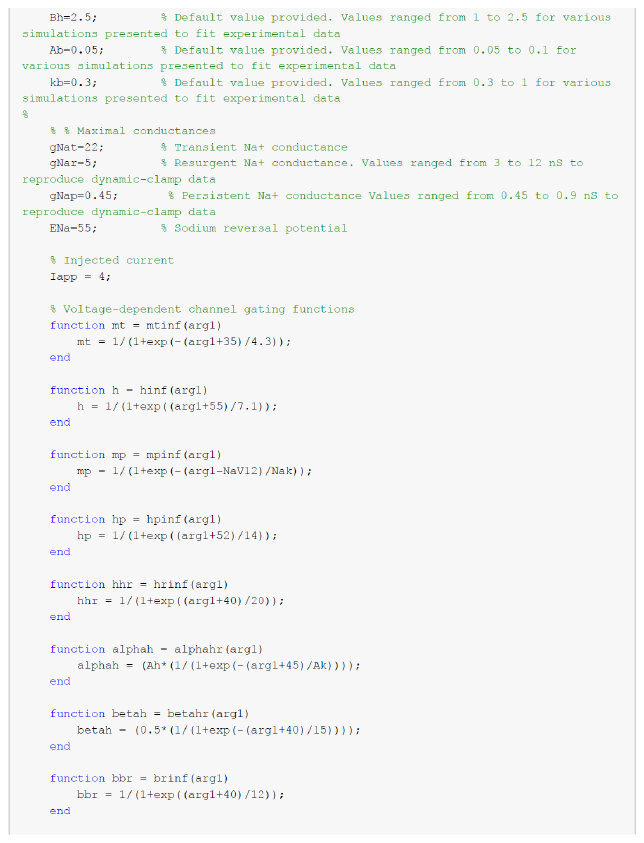

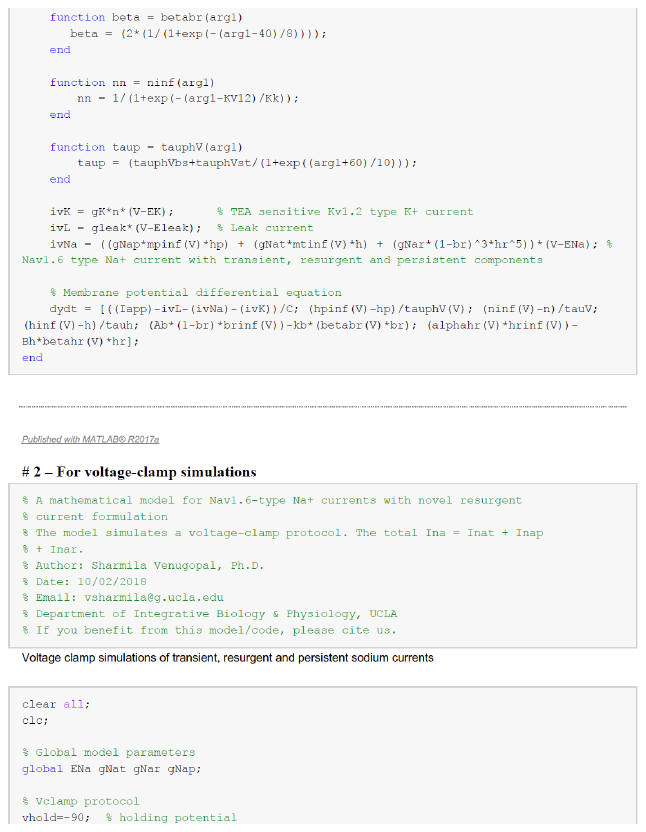

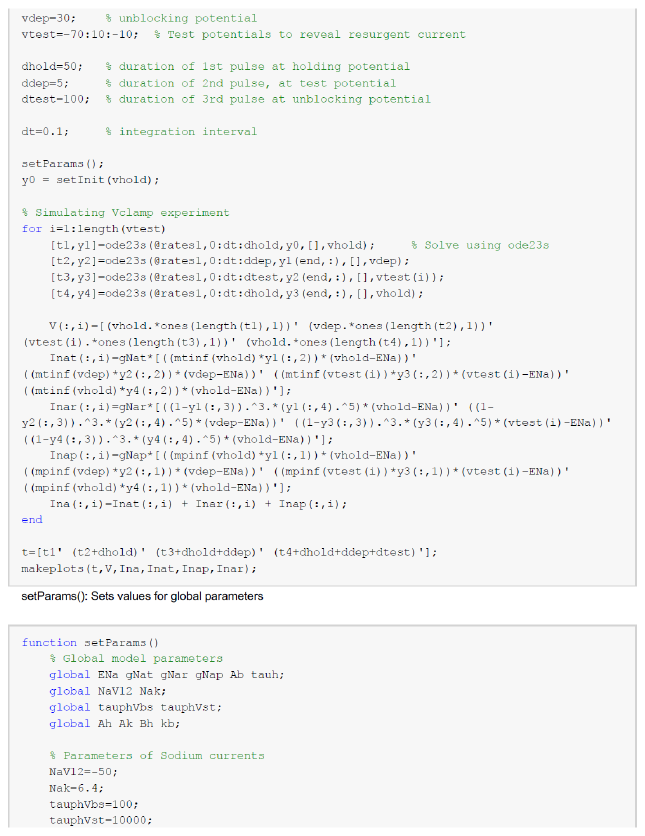

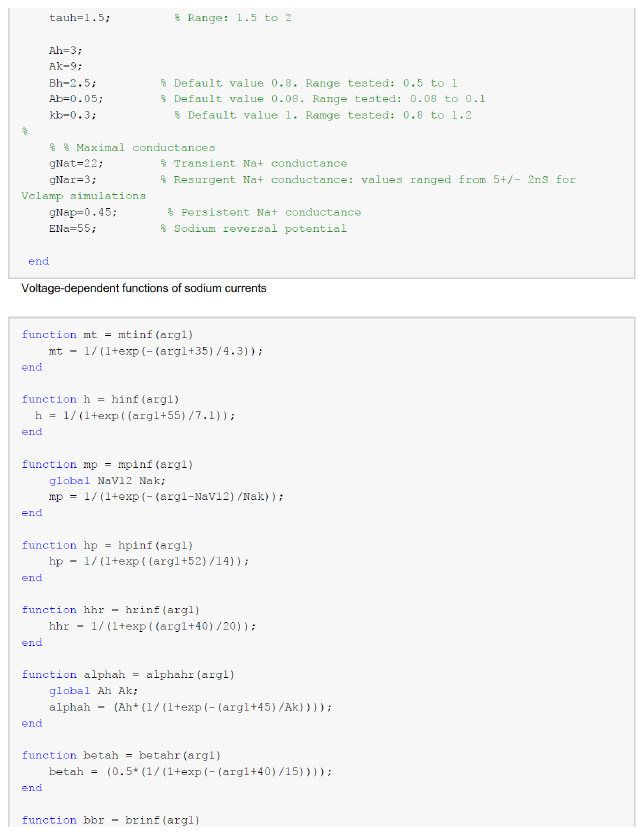

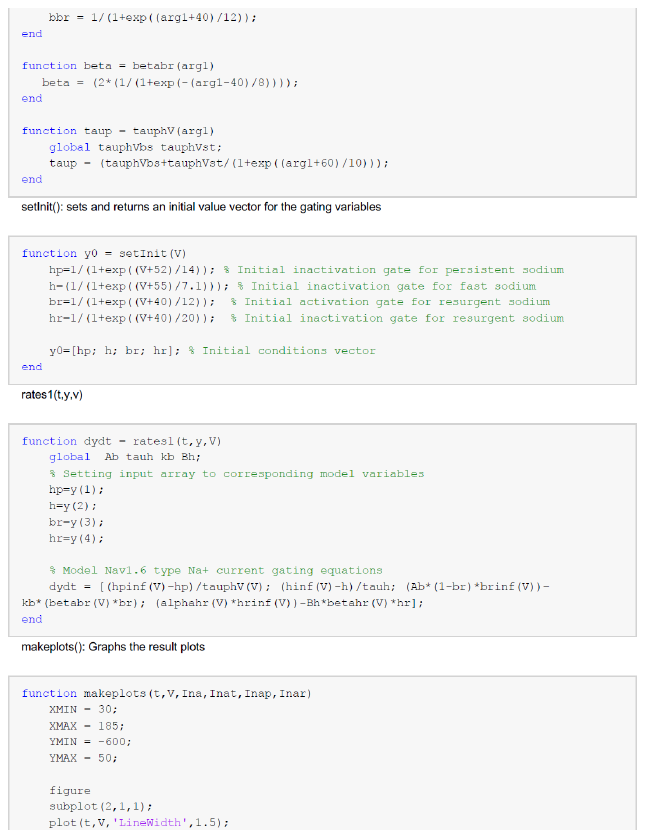

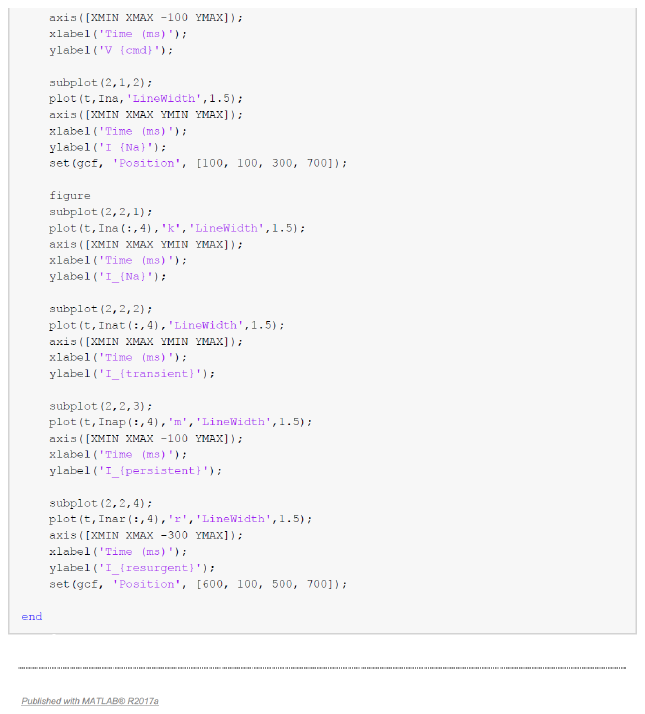

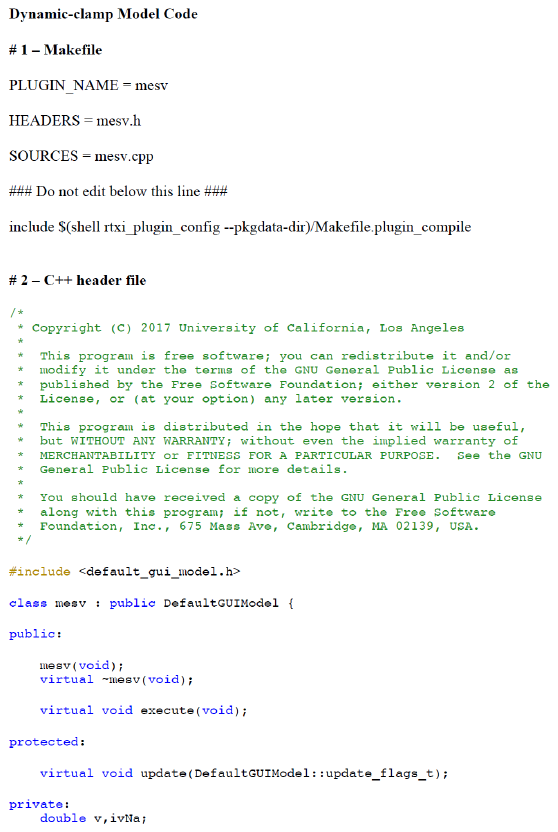

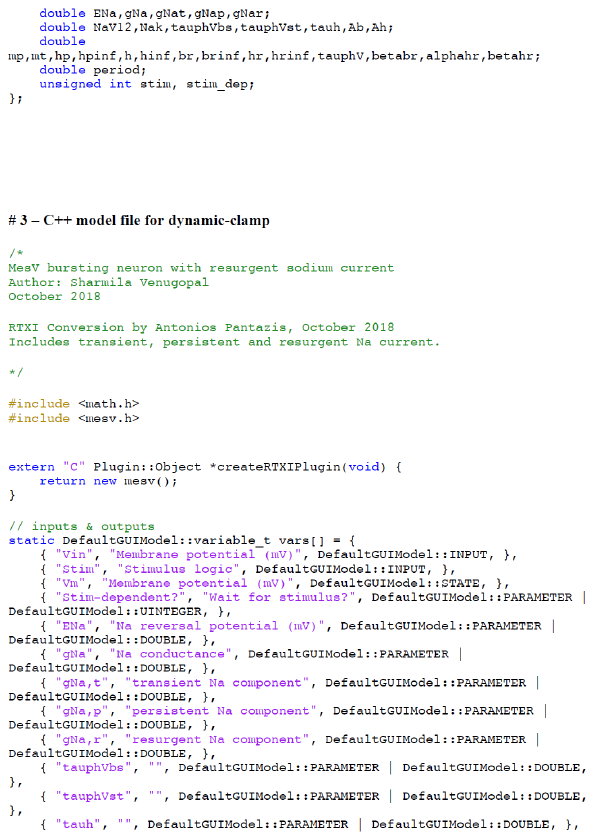

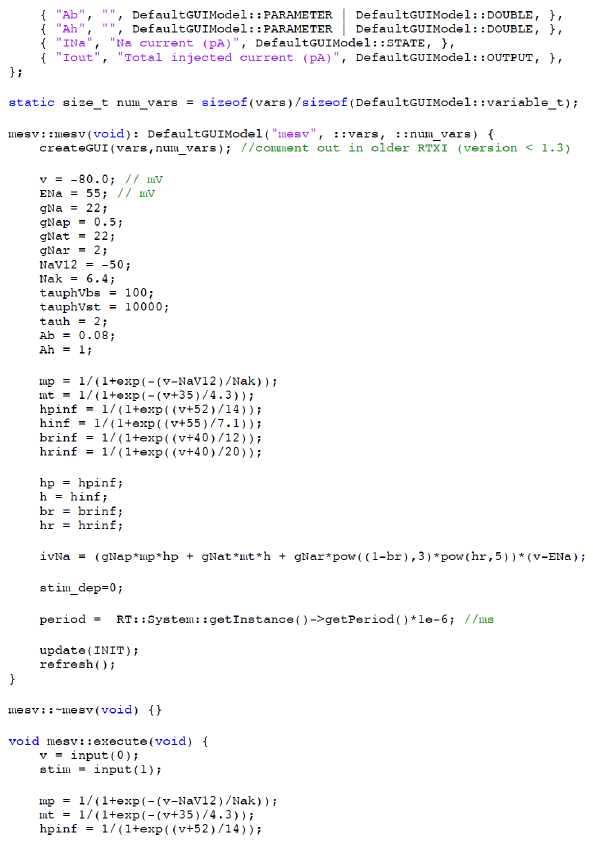

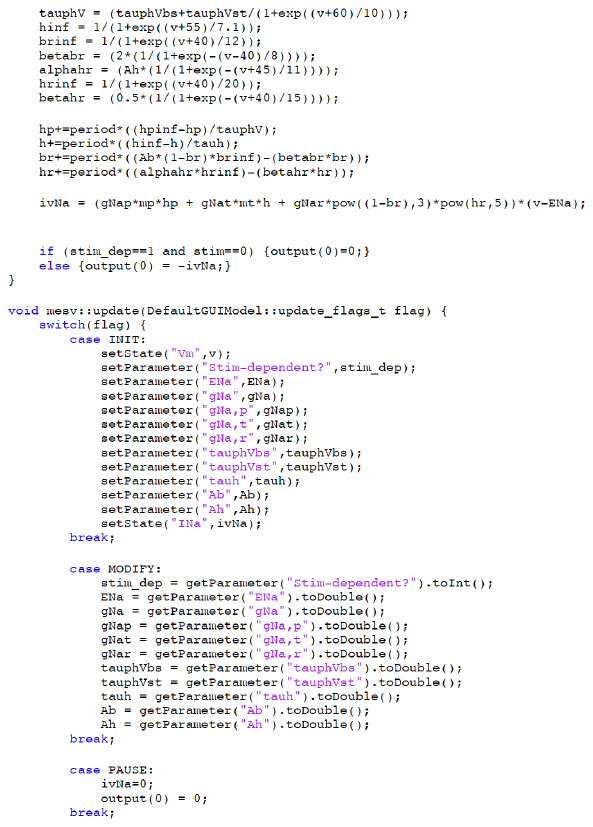

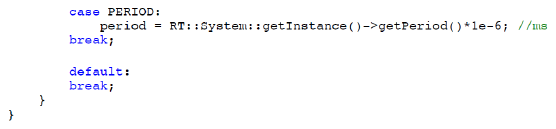
